# Evolutionary origins of protein novelty across an entire yeast subphylum

**DOI:** 10.64898/2026.08.29.748005

**Authors:** Emilios Tassios, Nikolaos Pirgelis, David Rinker, Eirini M. Tzermpou, Chris Todd Hittinger, Antonis Rokas, Christoforos Nikolaou, Nikolaos Vakirlis

## Abstract

Novel protein-coding sequences fuel molecular and cellular evolutionary innovations and frequently contribute to species-defining characteristics. They can originate either de novo from previously noncoding sequences or through extreme divergence of already coding ones. How frequently each mechanism occurs and how they shape the structural and functional potential of the resulting proteins remains unclear. Here, we conducted a broad computational investigation of genetic and protein novelty throughout the entire subphylum of Saccharomycotina yeasts. We detected more than 5,000 robust de novo genes across 332 species and compared them to more than 6,000 novel genes resulting from extreme sequence divergence, revealing two quantitatively similar but qualitatively distinct modes of evolution of novelty. A remarkable 40% of de novo proteins are predicted to localize to mitochondria compared to only 20% of divergent, with the latter also being substantially longer and more disordered. A detailed analysis of conservatively predicted tertiary structures of novel proteins shows that “invention” of new folds occurs more frequently through de novo emergence. We also illustrate cases of evolutionary “re-invention” of existing protein folds from noncoding sequences. Our work deepens our understanding of the origins and importance of novel proteins, opening new directions for further structural and functional characterization.

## Introduction

What makes a species unique is a profound and consequential biological question. Part of the answer lies in protein-coding genes that are present in an organism but absent from most or all other species. Such novel genes (aka orphan genes or Taxonomically Restricted Genes, TRGs) have been an enduring genomic and evolutionary mystery since they were first discovered in the yeast genome ∼30 years ago^1^. But with the availability of massive amounts of genomic data, technological and conceptual advances, such novel genes are no longer quite as mysterious and there are currently two main explanations for their evolutionary origins.

The first is divergence. Novel genes can result through the accumulation of mutations, causing homologous sequences to lose all detectable similarity^2^. This process, which may or may not be accelerated by relaxed negative selection or positive selection^3,4^, is only now starting to be investigated systematically, but it is clear that sequence divergence beyond recognition represents a major process leading to novel genes at different taxonomic levels^2,5–8^. Yet, even when coupled to coding sequence remodeling, truncation or alternative reading frame utilization divergence cannot, it would seem, account for the entirety of novel genes^8^, especially species-specific ones which frequently reach up to 30% of a species’ gene catalogue^9,10^, with even higher values in some viral genomes^11,12^.

The flipside to divergence is de novo gene birth, the evolution of an entirely new protein-coding gene out of a previously noncoding region of the genome^13^. De novo genes have been linked to phenotypic and morphological novelties^14–17^ and over the past two decades the process of de novo gene birth has grown from controversial to firmly established. Still, identifying bona fide de novo genes remains a challenging task involving trade-offs between confidence that a sequence is functionally protein-coding and that it originated de novo.

Yeasts, particularly *S. cerevisiae*, have served as an invaluable model in the study of de novo gene birth since the very beginning^18,19^. Past studies have offered strong support for the prevalence of robust bona fide de novo genes across multiple *Saccharomyces* and *Lachancea* yeast genomes and the frequent expression of incipient proto-genes that are likely evolutionarily short lived^20–23^. *S. cerevisiae* has also served as a model for understanding the structural and functional potential of intergenic Open Reading Frames (ORFs) that might one day evolve to be protein-coding^24–26^. Basic protein structural elements, such as α-helices, can be readily obtained from yeast intergenic ORFs^25^, and across species such sequences are predicted to be mostly foldable^26^. The underlying genomic sequences bias the properties of such elements^27^: in *S. cerevisiae*, intergenic ORFs have an even higher than expected potential to form transmembrane (TM) domains^28^, a property shared by almost all Saccharomycotina genomes^24^. Recently we discovered that this tendency is explained by polyA/T tracts, which are thought to primarily act as nucleosome depletion motifs and are abundant within the *S. cerevisiae* genome^29^, though not present in all yeasts^30^.

The prevalence of TM domains in young *S. cerevisiae* de novo proteins has recently been linked to localization at the endoplasmic reticulum (ER) and degradation by a common pathway^31^. Other targeted studies in model species like fruit fly, mouse, rice, and others have revealed functions for specific de novo proteins (reviewed in refs^32,33^) and a tendency towards reproduction-related activities. So far, high-throughput functional characterization of de novo protein function has not been feasible, but the advent of tools harnessing protein Language Model (pLM) embeddings like DeepFRI^34^, GoPredSim^35,36^ or PSALM^33^ has opened the way for at least tentative estimations of functional trends in the protein dark matter^37^.

The structural properties and novelty of de novo proteins is another critical question which can now be addressed with structure prediction tools like AlphaFold2^38^ and ESMfold^39^. Do de novo proteins tend to assume novel or existing folds? If so, is this due to or despite biases in composition, repeats or other motifs in noncoding regions? This question pertains to how we view and study extant novelty, but it is also a means to understand how the original “invention” of basic protein folds could have unfolded early in life’s history^40–42^. One hypothesis is that repetition of simpler elements is a dominant mechanism in the evolution of folds, which is supported by the fact that some of the most common folds are repetitive^43,44^. A few studies on de novo protein datasets in Drosophila^45–48^ have shown that they can assume both novel and existing folds, but this remains a mostly open question. Additionally, the suitability of the new generation of structure prediction methods for short, entirely novel protein sequences, such as those originating de novo, is unclear^49–51^ as methods find it challenging to confidently predict a structure for such sequences. With accumulating evidence showing that many novel structures and even basic folds remain to be discovered^52–54^, elucidating their evolutionary sources becomes more salient than ever. To date few studies have attempted large-scale surveys of the origins of novel genes and none has covered more than a few tens of species^20,21,55–57^. Here, we present a broad-scale computational investigation of novel genes and their evolutionary origins covering an entire subphylum, the Saccharomycotina yeasts. Building on a large, dense genomic dataset, we define a catalogue of de novo and diverged novel genes supported by multiple lines of evidence. Through this approach emerges a juxtaposition of de novo gene birth and extreme sequence divergence, as two distinct modes of evolution of novelty, resulting in proteins with significantly different characteristics. We also provide evidence for the de novo “re-invention” of both simple and more complex protein folds from noncoding DNA.

## Results

### A comprehensive catalogue of Saccharomycotina-specific genes and their long-term evolution

To study the origin and evolution of novel genes, we first sought to identify Taxonomically Restricted Genes (TRGs) within Saccharomycotina. Starting with the complete catalogue of Saccharomycotina proteins clustered into 233,478 homologous families by Shen *et al.*^58^ together with proteins from 11 outgroups^58^, we excluded 47,357 clusters containing at least one outgroup gene, as these have originated before the divergence of the clade and a further 36,157 families containing at least one protein with sequence similarity to proteins in other outgroups (see **Methods** and **Figure 1**). This resulted in 388,872 genes, organized in 145,417 TRG families, of which ∼70% were singletons. The phylogenetic branch of origin and thus age of each family was estimated by taking the most recent common ancestor of all species with a homologue (Dollo parsimony; see **Methods**). To ensure that we were not underestimating the phylogenetic age of any family, we performed an additional round of clustering (hereafter re-clustering) of the original families based on more relaxed similarity criteria (see Supp. Figure 1A,B and **Methods**). This resulted in 100,249 final TRG families consisting largely of singletons with a recent branch of origin (see distribution in Supp. Figure 1B).

**Figure 1.**
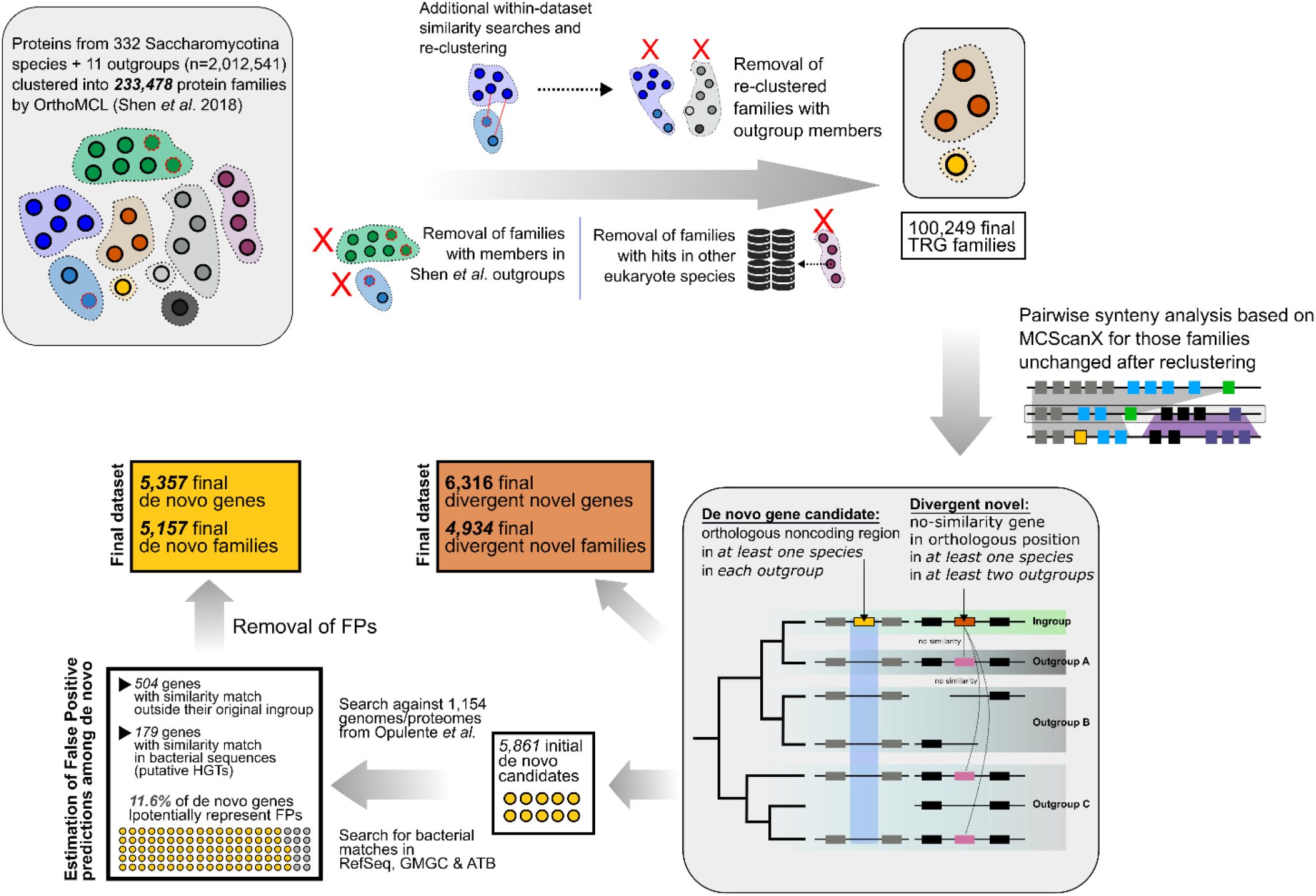
Graphical summary of the main steps of the pipeline for the identification of Taxonomically Restricted Genes (TRGs), divergent novel genes, de novo genes, and estimation of false positives among the latter. See main text for details. Synteny analysis shows idealized cases.

We next examined long-term trends of gene and protein properties over time. First, we counted the number of final TRG families originating at each branch of the phylogenetic tree (number of origination events; root branch excluded) and found a significant correlation to branch length (Pearson’s r=0.503, P=2.27e-36; Supp. Figure 2). This suggests that novelty originates at a steady evolutionary rate.

We then plotted averages of gene and protein properties binned according to their predicted age, for species belonging to each of the 20 Saccharomycotina genera with at least 5 species (Supp. Figures 1, 3, 4 and 5). We plotted trends using ages based on 1) the original clustering, which tends to split larger homologous families into smaller orthologous groups, and 2) our re-clustered final families, which tend to group homologous families (see Supp. Figure 1A). This allows us to analyze two versions of evolutionary trends: those reflecting processes, such as duplication and divergence (and other events such as gene fusions etc.), and those that only reflect true gene family founder events. Are there crucial differences between “true” emergence of sequence novelty versus diversification of existing families?

As expected given previous findings^59^, age correlates with length: younger genes are shorter than older genes, but the trend is less steep in the re-clustered version (Supp. Figure 1C). In contrast, GC content is remarkably unchanged in almost all genera in both versions, revealing important stability in the nucleotide composition despite significant changes in higher level properties (Supp. Figure 1C). Various structural properties correlate to GC content, including intrinsic structural disorder (ISD)^13,60^. Under the original clustering, we observed a strong trend of decreasing average ISD with evolutionary age (Supp. Figure 1C). This trend was weaker in the re-clustered version but young proteins were still more disordered than ancient ones, suggesting that the evolutionary processes that give rise to novel proteins bias them towards disorder. We found no trends in propensity to form transmembrane (TM) domains (Supp. Figure 3A) or biosynthetic cost (Supp. Figure 3B). Additionally, slight trends in protein secondary structure under the original clustering disappear when re-clustering (Supp. Figure 4) as do trends in percentage of residues predicted to be exposed to solvent (Supp. Figure 5). Overall, the weakening of trends after re-clustering can be explained by initially young proteins moving to the conserved bin, suggesting that these trends are partly driven by strongly divergent families. (i.e. split in the original clustering but merged during re-clustering)

#### More than 5,000 high-confidence de novo genes in Saccharomycotina

To distinguish those families that have most likely originated de novo from previously noncoding DNA, we applied a stringent synteny-based pipeline^61^ (see **Methods** and **Figure 1**). For each TRG family, we identified regions of conserved microsynteny in the three closest outgroup phylogenetic clades defined by its branch of origin and kept cases without annotated protein-coding genes in the orthologous regions. This resulted in an initial set of de novo gene candidates, of which 5,540 were species-specific and 321 belonged to multi-species families. As expected, we observed a negative correlation between the number of de novo genes present in each species and the age of the furthest species used as an outgroup (Spearman’s ρ = -0.38, P=1.23e-11; Supp. Figure 6), establishing that phylogenetic distance limits detection.

Yeasts are known to exchange genes with bacteria^62^, which raises the possibility that some of our de novo gene candidates could actually represent Horizontal Gene Transfers (HGTs). To identify such cases, we performed three similarity searches: 1) against the entire bacterial and archaeal RefSeq protein catalogue downloaded from NCBI, 2) against the Global Microbial Gene Catalogue database (GMGC) and 3) against the bacterial genomes found in AllTheBacteria (ATB)^63^, a collection of over 2 million bacterial genome assemblies. In total, 125 proteins had a significant protein-level match and an additional 60 mapped to at least one ATB genome of high quality (see **Methods;** Supp. Table 1). These 179 de novo candidates could be novel HGTs in either direction but could also represent contaminations on both the yeast and bacterial sides. We nonetheless decided to filter them out as potential false positives and leave their full analysis for future work.

While this work was in preparation, a richer dataset of Saccharomycotina genomes was published^64^. This presented a unique opportunity to benchmark our approach for identifying de novo genes by quantifying false positives: how many of the initial de novo candidates find homologues in outgroup species once taxonomic sampling is increased? Using protein-level similarity searches and protein-to-genome mappings, we identified false positives 504 (FPs) (8.5%). These were then split into two groups: those where the extra species increased the estimated timing of origination less than 20 my and those where it increased more than 20my (Supp. Figure 7A). Both FP groups were substantially longer than the rest (Supp. Figure 7B), in line with them being older than initially estimated. Additionally, both groups of FPs had more hits than the remaining de novo genes (Supp. Figure 7C). Again, this suggests that these are indeed older genes that should not be classified as de novo. Overall, then, by increasing the target dataset ∼three-fold, we obtained an upper limit of false positives of 11.6%, including the HGT cases. These were removed, and thus if any false positives remain, they are expected to be lower still.

The final number of de novo genes is 5,357 (Supp. Table 2), of which 5,127 are species-specific and 230 are multi-species (5,157 families; **Figure 2A**). An example of a de novo gene in the species *Hyphopichia heimii* aligned to its orthologous region in outgroups is shown in **Figure 2B**. De novo genes are ubiquitous: 91.2% (303/332) of Saccharomycotina species have at least one de novo gene with an average of almost 17 genes per genome (**Figure 2C**). As expected, species at tips of longer branches have fewer de novo genes due to synteny limitations yet some shorter branches seem to have higher rates of de novo origination than others (darker branches in **Figure 2C**). De novo genes are known and indeed expected to be short, and we confirm this here, with species-specific de novo genes across Saccharomycotina orders being ∼4 times shorter than conserved genes (**Figure 2D**) and multi-species ones being slightly longer (species-specific: 300.4nt, multi-species: 336.4nt), likely reflecting their elongation and increase in complexity over time. We found low correlation between the number of de novo genes and the number of BUSCO genes (Supp. Figure 8A) and no correlation to assembly N50 (Supp. Figure 8B), showing that assembly quality is not affecting our analyses.

**Figure 2.**
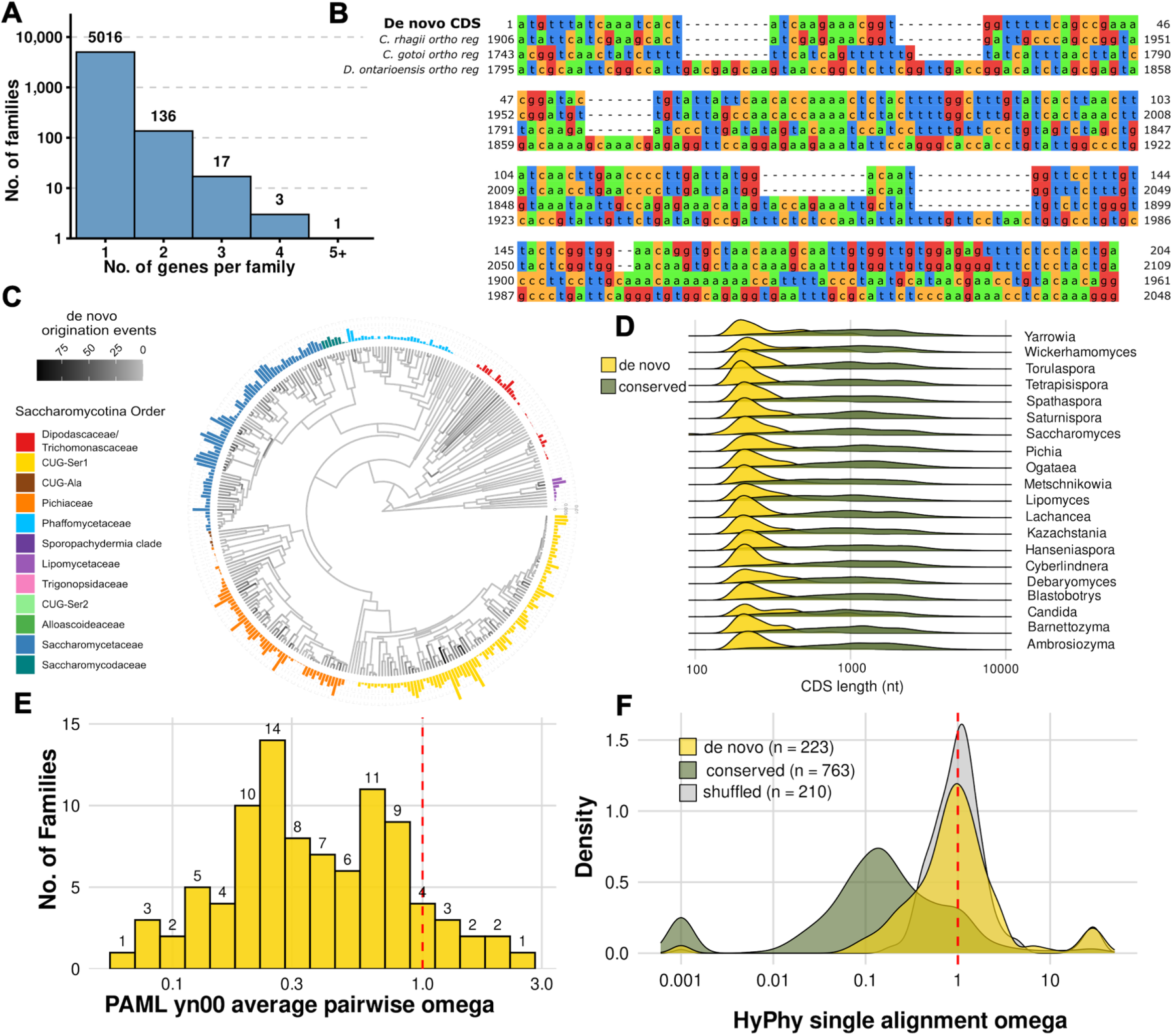
**A**) Distribution of sizes of de novo gene families. **B**) Alignment of a de novo gene’s CDS (*Hyphopichia_heimii@Seq_4498)* to the orthologous regions of its three outgroups. **C**) Distribution of percentages of de novo genes over all genes in the genome across different species and number of de novo origination events along the branches. **D**) CDS length comparison between de novo and conserved genes. **E**) Distribution of *d_N_*/*d_S_* values of 92 multispecies de novo gene families, as calculated using yn00 of PAML. **F**) Distribution of omega values (HyPhy) calculated from intra-specific alignments of de novo and conserved genes. Shuffled alignments of de novo genes are shown as negative controls.

Because our work is based on ab initio gene predictions, we also define an even stricter subset of de novo genes that 1) have only 1 exon, and 2) can be fully mapped onto the assemblies of Opulente *et al*., when the genome was part of that dataset as well. This subset, denoted “strict de novo”, consists of 2,590 genes (Supp. Table 3).

Short genes that lack homologues carry a non-negligible risk of being spurious annotation artefacts in which case they would lack signatures of natural selection. This, however is not the case for de novo genes: we measured selection by calculating the ratio of non-synonymous to synonymous mutation rates (*d_N_/d_S_*) on those present in at least two species that resulted in suitable alignments (n=92; see **Methods**) and found clear evidence of purifying selection (mean = 0.51) which supports their functional protein-coding status (**Figure 2E**). When using only strict de novo genes (n=62), our observations were similar (mean=0.47) and were not statistically different (W=3016.5, P=0.72) (Supp. Figure 9A).

We next applied a similar approach for species-specific de novo genes based on intra-specific alignments; aware of its caveats^65,66^. Young genes have been shown by several studies to be either under weak selection pressure or even under positive selection, independently of their essentiality^13,21,56,67–70^. To understand how to interpret these intra-specific selection estimates, we first calculated omega values from intra-specific alignments of conserved genes (see **Methods**). These were mostly found to be under purifying selection (mean 0.76 and median 0.14 respectively; **Figure 2F**), while their distribution exhibited two clear peaks, one at ∼0.5 and one at 1. The distribution of values for de novo genes had its main peak at ∼1, but a clear secondary peak matching the main one of conserved genes. In contrast, shuffled alignments of de novo genes centered sharply around 1. Given that this approach is sensitive enough to only pick up signal from genes under strong purifying selection^65,66^, our results show that at least a subset of species-specific de novo genes is under strong selection, a smaller subset than that of conserved genes perhaps, consistent with previous studies. The distribution was similar for the strict de novo dataset (Supp. Figure 9B). Overall, we provide strong support for both the de novo status and the functional, protein-coding status of de novo genes.

### Sequence divergence beyond recognition as a distinct process of novelty

Novel genes can also result from extreme sequence divergence beyond recognition and the importance of this process for evolutionary and functional protein novelty is understudied. We thus sought to identify such diverged novel genes and compare them to de novo originated ones. We identified “invisible” homologues of TRGs using similar synteny criteria as for de novo genes and retained cases where a gene belonging to a different family is found in the orthologous syntenic region of at least one genome in at least two of the three outgroups (see **Figure 1** and **Methods)**. Additional similarity filters were then applied to obtain a set of divergent novel genes. The lengths of divergent novel genes and their “invisible” homologues correlate (r=0.41, P<2.2e-16), supporting their status as homologues (**Figure 3A**). In total, we identified 6,316 divergent novel genes (Supp. Table 2). They are clustered into 4,934 families (**Figure 3B**) and distributed across 326/332 species (**Figure 3C**). There is good correlation between the number of events of de novo and divergent origination (Pearson’s r=0.580, P=6.26e-46; **Figure 3D**), partly explained by branch length (i.e. longer branches tend to have both more divergent and de novo events); indeed, when controlling for branch length the correlation drops substantially (Spearman’s ρ = 0.38, P=1.58e-18; Supp. Figure 9). The relatively close overall numbers of divergent and de novo genes suggest that the contributions of the two processes to novelty are quantitatively comparable.

**Figure 3.**
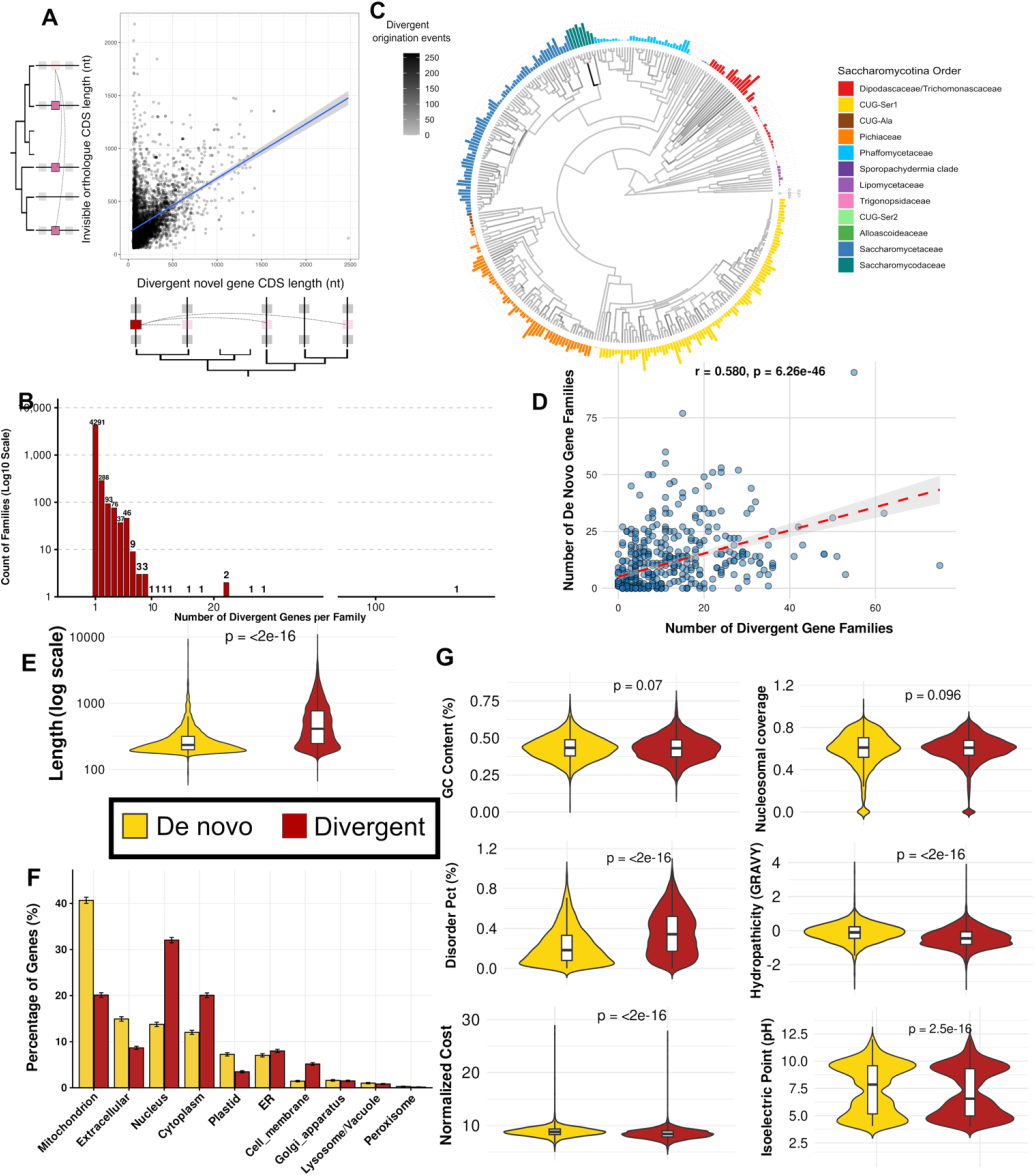
**A)** Scatterplot of CDS lengths of divergent novel genes (n=5,469; X axis) to the average CDS length of their “invisible” homologues (Y axis). **B**) Size distribution of divergent novel gene families. **C**) Proportion of divergent genes per species (bars) and numbers of divergent origination events per branch. **D**) Correlation of the number of originations events of de novo and divergent families. **E**) CDS length distributions of de novo and divergent genes. **F)** Subcellular localization predictions of de novo and divergent proteins using DeepLoc1. **G**) Comparison of gene and protein properties between de novo and divergent groups. Wilcoxon test P-values are shown.

We next hypothesized that, if de novo genes really have their origins in noncoding sequences, while divergent genes have their origins in coding sequences, we should expect them to differ in sequence and biophysical properties. Indeed, we found that de novo genes are significantly shorter than diverged (**Figure 3E**), even when only considering recently originated ones thereby controlling for the fact that diverged are on average older than de novo (Supp. Figure 10). This can be explained by the mechanism of origination itself, as sequence divergent genes have their evolutionary origins on longer coding sequences, whilst de novo genes must evolve from necessarily small genomic Open Reading Frames (ORFs).

De novo proteins show a striking subcellular localization enrichment (predicted by DeepLoc^71^), with 40.6% of them predicted to localize to the mitochondria (**Figure 3F**; proportion in strict dataset is 43.2%) compared to only 20.1% in divergent. To control for length differences, we also predicted the localization of a shuffled version of the de novo proteins and randomly drawn segments of conserved proteins that had the same length distribution as the de novo ones. Both sets had lower percentages of predicted mitochondrial localization than de novo genes, 30% for the shuffled and 21% for the conserved control (Supp. Figure 11, Supp. Table 4). This indicates that both amino acid composition and length are factors but do not fully explain the enrichment seen in de novo proteins. This substantial proportion over such a large dataset strongly supports a significant functional role of small proteins of *specifically de novo origin* as mitochondrially functioning peptides, or mito-SEPs^72,73^. De novo proteins are also strongly enriched in extracellular localization (16.7% vs. 8% in divergent).

Diverged proteins are significantly more disordered than de novo ones (mean 35.7% and 22.8% respectively, P < 2.2e-16), a difference that persists when correcting for age and across most yeast genera (**Figure 3G**, Supp. Figure 10,12). This is consistent with the fact that disordered proteins evolve much faster than globular ones as they are more flexible and can accommodate extensive sequence variability^74,75^. De novo proteins are more hydrophobic and have higher predicted biosynthetic cost, consistent with them being less optimized than divergent ones (**Figure 3G**, both P<2.2e-16, both true when correcting for age; see Supp. Figure 10). Previous works have suggested that putative de novo proteins have higher isoelectric point (pI) relative to conserved ones^20,23^; we find that in our dataset they have higher average values than divergent ones, including across most yeast genera when analyzed separately (Supp. Figure 13) and indeed higher than conserved proteins as well (Supp. Figure 14). Overall, these differences provide consistent, solid support for two distinct evolutionary processes, producing novelty of two different “flavors”.

### Repetitive intergenic motifs corroborate distinct evolutionary origins for de novo and divergent novel genes

Biases and patterns found within intergenic genomic regions may influence the functional and evolutionary trajectory of de novo genes and the proteins they encode. We previously showed that in the *S. cerevisiae* genome, motifs made of tracts of consecutive adenines (polyA), thymines (polyT; together, polyA/T) and tracts of TA repeats (TATA), which are enriched and abundant in intergenic regions, are co-opted by emerging de novo genes as sequences encoding predicted transmembrane domains^29^ (motif logos shown in **Figure 4A**). They are ultimately responsible for the enrichment of transmembrane domains in de novo proteins in *S. cerevisiae*, and they also serve as an important signature of intergenic regions as they are exceedingly rare in conserved protein-coding genes. Here we expanded the search for these specific motifs to the entire subphylum.

**Figure 4.**
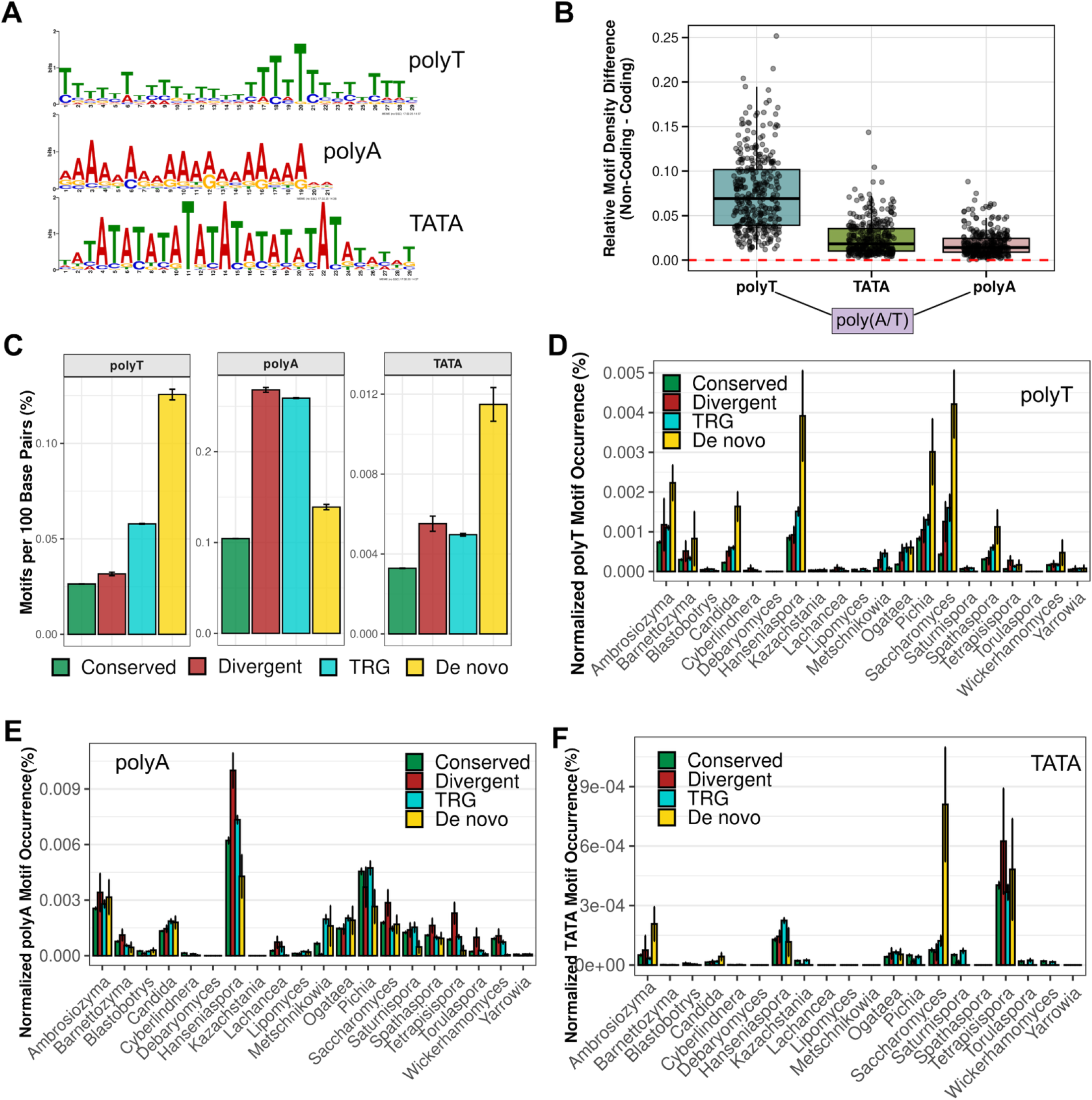
**A**) Sequence logos generated by MEME for the three motifs identified by Vakirlis and Fuqua in the *S. cerevisiae* genome. **B**) Relative difference of motif density between non-coding and coding regions for 332 genomes, for the three motifs shown in A. Only forward strand is searched and thus polyT and polyA distributions should be viewed as one, corresponding largely to the same sequences. **C**) Motif occurrence per 100 base pairs for the three motifs in conserved, divergent, TRGs and de novo gene CDSs. **D**) Same as C but only for the polyT motif across genera with at least 5 species. **E**) Same as D but for the polyA motif. **F**) Same as C but for the TATA motif.

First, we asked whether these motifs characterize intergenic regions in all species as they do in *S. cerevisiae*. We found that in all 332 species the polyT motif is substantially more prevalent in noncoding regions compared to coding regions (**Figure 4B**), as is the TATA motif but to a lower degree. Note that we only search the forward strand and so the enrichment for polyA and polyT should be viewed jointly, as one, since they are each other’s reverse complement. Thus, polyA/T tracts (and secondarily TATA) are a reliable intergenic marker, and as such, we can expect to find it more frequently in genes of de novo origin compared to the rest. Indeed, this is what we observe: the polyT motif specifically is found 5.96 times more in de novo genes’ CDSs than conserved genes and 3.4 times more than divergent ones (**Figure 4C**; note that the difference in GC% between de novo and divergent is statistically insignificant, see **Figure 3G**). This enrichment holds when analyzing separately different genera: especially in those genera where polyA/T tracts are more abundant, de novo genes always have significantly more polyT motifs than genes in other groups **(Figure 4D**). Conversely, across genera, polyA tracts are consistently equally or more abundant in divergent novel genes and TRGs than de novo ones (**Figure 4E**), while TATA motifs vary significantly from genus to genus (**Figure 4F**). Thus, analysis of these motifs strongly supports the distinct evolutionary origins of de novo and divergent novel genes and the preferential evolution of de novo genes on polyT tracts.

### Novel and existing structural folds evolve both de novo and through divergence

How often does sequence novelty, evolved de novo or through divergence, translate to structural novelty? To address this question, we first used the ESM3 protein language model, which has been shown to capture protein structure well^76^. We generated ESM3 embeddings of all species-specific de novo, divergent and a sample of conserved proteins and plotted them using UMAP (**Figure 5A**; only species-specific de novo and divergent are shown to sharpen the comparison to conserved). Within this embedding space, de novo proteins seem to be restricted to specific areas, relative to both conserved and divergent ones, providing evidence for an emergence bias^27^. We next predicted tertiary protein structures using ESM3 for all de novo and divergent novel proteins, as well as entire intergenic regions where in-frame stop codons have been removed (iORFs from ref^24^; see **Methods**). As controls, we also predicted structures based on shuffled versions of the original sequences in these three groups as well as random protein sequences drawn from a uniform amino acid distribution. Shuffled sequences, as well as translated iORFs, showed comparable distributions of predicted structural confidence metrics to real sequences of de novo and divergent novel proteins (predicted Template Modeling, pTM, in **Figure 5B**; average predicted Local Distance Difference Test, pLDDT in Figure 15A). Even completely random proteins produced comparable metrics. Structures of divergent novel proteins had significantly lower confidence, due in part to them being longer and more disorder as both protein length and ISD correlate negatively to pTM (length: r =-0.39; ISD: r=-0.54, both P < 2.2e-16).

**Figure 5.**
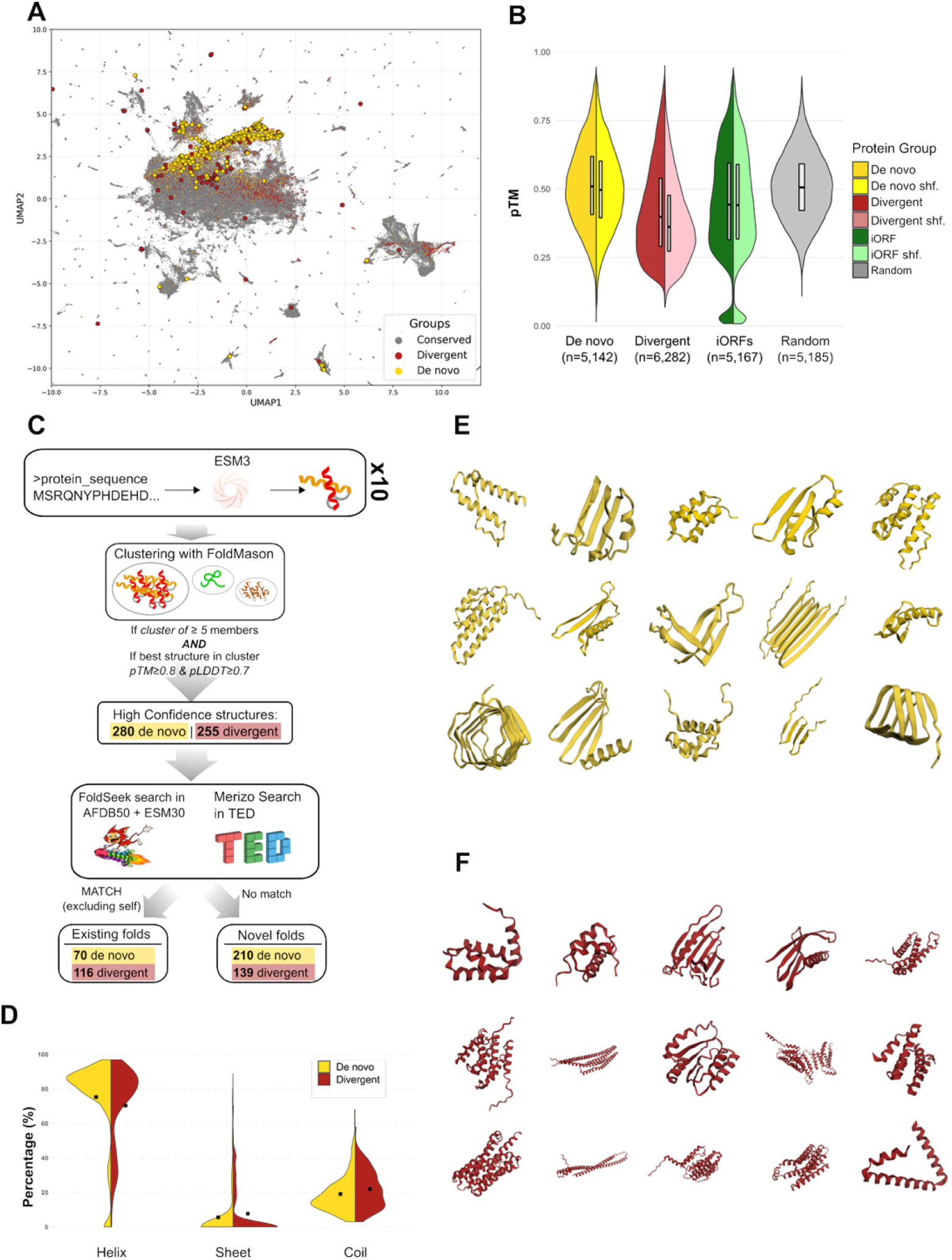
**A)** UMAP projection of ESM3 embeddings of all species-specific de novo (n=5,127) and divergent (n= 4,396) proteins and a sample of conserved Saccharomycotina proteins with origin at least as old as the root (n= 286,243). Points with black contours correspond to proteins that have produced high-confidence novel structures. **B**) Distributions of pTM values for original proteins and matched shuffled controls, as well as a sample of random proteins with uniform amino acid distribution and matched lengths to the de novo set. **C**) Main steps of the strategy used for structure prediction and similarity search. **D**) Secondary structure element percentage distributions for de novo and divergent high confidence structures as predicted by ESM3 based on the tertiary structure models. **E**) Examples of ESM3 predicted novel structures of de novo proteins. **F**) Examples of ESM3 predicted novel structures of divergent proteins.

Given that predicted confidence metrics were high even for randomized sequences, to obtain robust structures we performed ten replicates and only kept those where a similar structure was predicted at least 5 times (see **Figure 5C** and **Methods**). Real sequences resulted in replicable structures more frequently than shuffled ones although the difference was much smaller for de novo than divergent (Supp. Figure 15B). Applying an additional strict confidence cut-off (pTM≥0.8 and pLDDT≥0.7), we obtained a total of 280 high confidence de novo and 255 divergent structures (Supp. Table 4), which we retained for downstream analyses. Secondary structural element analysis of predicted models revealed moderate but significant differences, with de novo proteins containing more helices, while both being overall helix-rich (**Figure 5D**). Note that in this and following structural analyses we necessarily do not include proteins with high disorder, which are more enriched in the divergent group, partly distinguishing it from de novo while likely also influencing its function.

Focusing on de novo first, we searched with FoldSeek for similarity to known structures within the structural databases AlphaFoldDB50 and ESM30 and found 16 de novo structures with matches (Supp. Table 5). Using Merizo^77,78^, we also detected protein domains and used them to search for similarity within The Encyclopedia of Domains (TED), finding an additional 54 (Supp. Figure 15C), for a total of 70/280 (24.5%; 68/277 unique families) (Supp. Table 6). In the divergent group this percentage is substantially higher at 40.3% (74 FoldSeek + 42 additional in TED out of a total of 255; 84/208 unique families, Supp. Table 5 and 6). This is expected if divergent novel protein sequences retain their ancestral structure despite drastic sequence changes. To confirm this, we searched for structural similarity between divergent novel proteins and their “invisible” homologues and found it for 21.4% of protein pairs (123/573), involving 39 out of 104 divergent proteins with high confidence structures and at least one invisible homologue (see **Methods**). For all of these 39 proteins, their structures also have at least one match to structures/domains in public databases and can thus be transitively inferred to have retained some similarity to their ancestral structural state. In sharp contrast, structural similarity only occurs for 0.9% of all other pairs (i.e. excluding pairs of divergent novel proteins and their “invisible” homologues; 279/30,315, involving 18 out of 104 divergent proteins). Overall, while both de novo proteins and divergent proteins have novel sequences, the latter adopt familiar folds much more frequently, in line with their predicted mode of evolution.

There are 210 de novo proteins (75%) and 139 divergent proteins (54%) with entirely novel structures, clustering into 207 and 135 structural families respectively (Supp. Table 7). Examples, found in **Figure 5E** and **F** exhibit plausible compact folds with some consisting of repeated units (all folds shown in Supp. Figure 16 and Supp. Figure 17). Like the rest of the high-confidence structures they are helix-rich (avg. 78.4%), with the divergent novel ones being significantly more helix-rich (80%) than divergent non-novel ones (58.6%). These findings strongly support the hypothesis that structural novelty is a frequent outcome of de novo emergence but can also result through divergence.

We next sought to identify cases of de novo “re-invention” of existing folds from non-coding regions as unambiguous as possible. To be maximally conservative, we focused on the strict dataset and also excluded those with predicted origin older than 50my, as these more ancient cases may yet be explained by a combination of divergence and gene loss (one more protein was removed; see **Methods**). Out of the remaining 26 (Supp. Table 8), we picked a few clear examples which can be seen in **Figure 6**. **Figure 6A** and **6B** show two representative examples of simple α-helix hairpin structures matching TED domains, both species-specific proteins with predicted mitochondrial localization. The alignments of the CDSs to their orthologous regions in outgroup species are consistent with absence of coding potential (multiple indels, low conservation, absence of ORFs), confirming their de novo status. **Figure 6C** shows an example of a highly repetitive de novo protein folding into a β-solenoid like structure, a domain known to be employed as scaffold for binding, adhesion, catalysis, secretion-associated virulence, and specialized mechanical or self-assembling roles^79,80^. This protein is variant-specific (*M. matae var. maris*) and so are most of the repeats, as can be seen in the orthologous region alignment. The 17aa-long repeat is also completely different to the one of the closest structural match (**Figure 6C**). These three cases exemplify de novo convergence of novel protein sequences towards both simple and repetitive structural domains. The fourth case (**Figure 6D**) presents an older and seemingly more mature de novo structure with two distinct identifiable domains matching an Aminohydrolase, N-terminal nucleophile (Ntn) domain (CATH superfamily: 3.60.20.10) and a novel TED domain identified on a protein from species *Meloidogyne enterolobii,* a root-knot nematode. This protein (Ambrosiozyma_pseudovanderkliftii@Seq_702) is part of a family of two with its origin at the ancestor of Ambrosiozyma (44.2mya) and the other member of the family also matches to a TED domain of the same CATH superfamily though it did not meet our similarity criteria to be included in this analysis.

**Figure 6.**
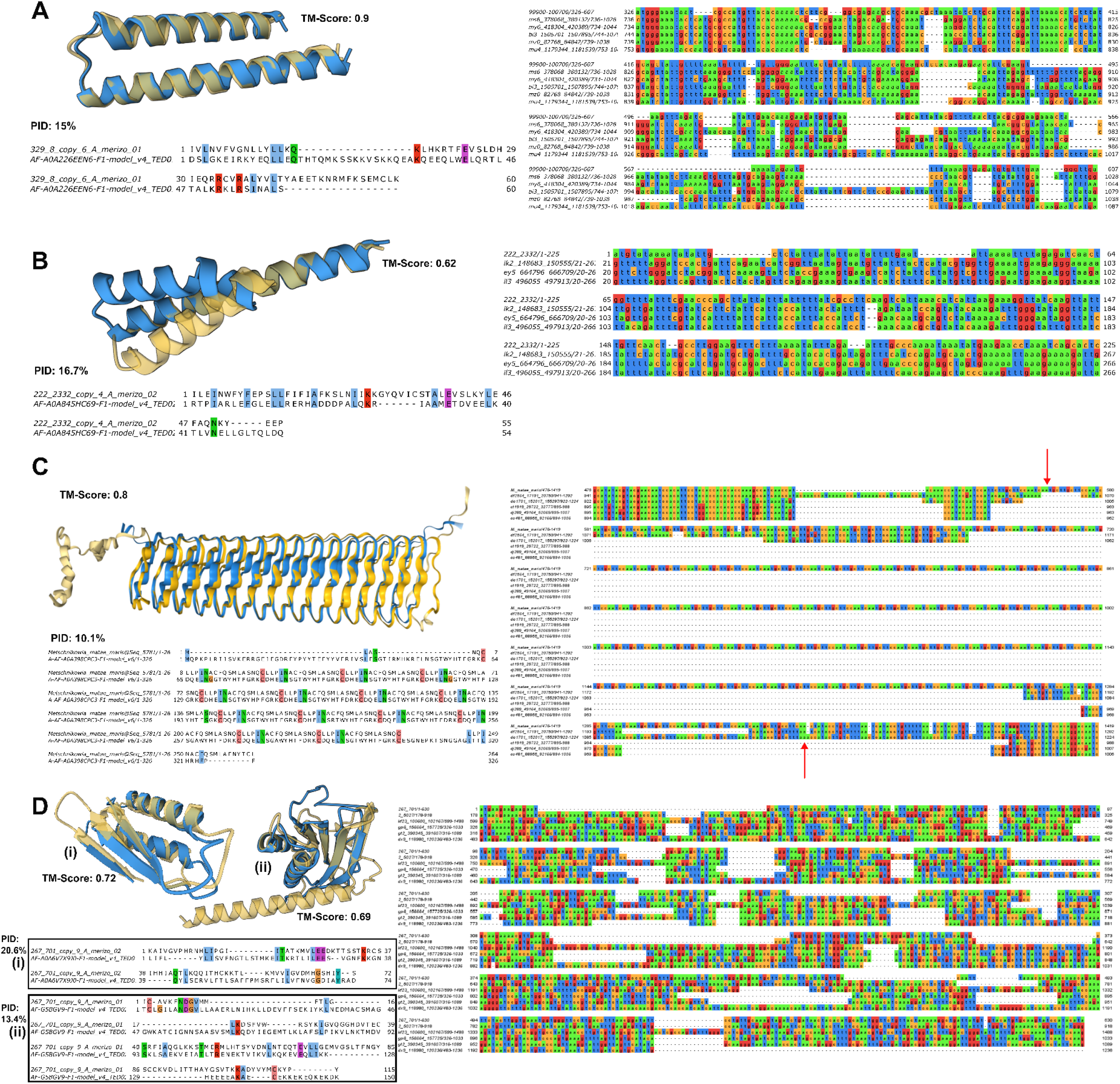
Structural and sequence alignments of de novo proteins to their corresponding best-matching domains/proteins structures and their sequences (left), and alignments of de novo CDSs to their orthologous regions in outgroup species (right; CDS is always the top sequences). De novo protein structures are shown in blue. **A)** Alignments for protein Debaryomyces_nepalensis@Seq_9: structural superposition to TED domain AF-A0A226EEN6-F1-model_v4_TED01 (top) and sequence alignment of the corresponding protein segment (bottom). On the right, MAFFT alignment to outgroup orthologous sequences. The sequence of the most distant outgroup has been removed for visibility as it contained large insertions. **B**) Same as in A for protein Tetrapisispora_namnaonensis@Seq_2333 and TED domain AF-A0A845HC69-F1-model_v4_TED02. **C**) Alignment of protein Metschnikowia_matae_maris@Seq_5781 to its best FoldSeek match from AFDB50, protein AF-A0A398CPC3-F1-model_v6. On the right, the orthologous region alignment is extended as the sequence consists almost entirely of repeats. Red arrows point to the start and stop codon positions of the CDS. **D**) Same as A and B for protein Ambrosiozyma_pseudovanderkliftii@Seq_702 and matches to two separate TED domains (i) AF-A0A6V7X9J0-F1-model_v4_TED01 and (ii) AF-G5BGV9-F1-model_v4_TED02. In the orthologous alignment, the second sequence from the top corresponds to its protein-coding homologue in *Ambrosiozyma monospora*.

To gain insight into the function of novel Saccharomycotina proteins, we used DeepFRI to predict Gene Ontology (GO) terms for all de novo and divergent proteins (Supp. Table 9). Significantly fewer de novo than divergent were assigned a GO term: 410/5,5357 de novo (7.6%) compared to 886/6,316 divergent (14%) but both proportions were low, so the main conclusion is that the vast majority of these novel proteins cannot be assigned a GO term even with cutting-edge predictive models. Nonetheless, those GO terms that do get assigned are strikingly similar between the two groups: the top three Cellular Component (CC) terms are for both groups “intrinsic component of the membrane”, “integral component of the membrane” and “non-membrane bounded organelle”; the top three Biological Process (BR) terms are for both groups “translation”, “peptide metabolic process” and “peptide biosynthetic process”; and the top three Molecular Function terms are for both groups “structural constituent of ribosome”, “structural molecule activity” and “RNA binding” (Supp. Figure 18). This leads us to hypothesize that despite their different routes of origination there is at least a small subset within each of these two groups that appears to function in membrane-associated translation and peptide biogenesis, potentially linked to the ER. This would be consistent with ER-related functions of de novo proteins in *S. cerevisiae*^81^.

Mitochondrial GO terms are virtually absent from this list (only 4 de novo genes match any) suggesting that DeepFRI captures a different biological aspect of our dataset, notably regarding subcellular localization. Indeed, those proteins with the top three membrane-related CC GO terms are strongly enriched in DeepLoc-predicted localization to the ER and this is true both for de novo (34%; 120/345) and divergent (26%; 200/756), whereas only 12% of these de novo localize to the mitochondrion (vs. a background of 40%, see Supp. Figure 18).

## Discussion

Using a broad and robust yeast genomic dataset, we were able to identify thousands of novel genes of de novo and divergent origin and to compare them at multiple levels and at an unprecedented phylogenetic scale. Building such a large set – to our knowledge the largest to date – allowed us to show that the two processes have a relatively similar quantitative impact within the pool of novel proteins. More crucially, it enabled us to uncover striking qualitative differences at sequence, structural, biochemical and functional levels which are most readily explained by the two distinct modes of origination. De novo proteins are shorter and less disordered than divergent, and they are also more positively charged. When exposed to an environment with pH equal to their isoelectric point, peptides show reduced solubility and become prone to forming protein aggregates^82,83^. With the cytoplasmic pH of the yeast cell being close to 7^84^, de novo peptides may be more susceptible to aggregate than divergent ones, perhaps a byproduct of their less optimized state. Their higher biosynthetic cost may be another such byproduct. De novo proteins exhibit markedly more structural novelty than divergent ones and, perhaps most importantly, at least four out of ten are predicted to localize and function at the mitochondrion, a proportion two times less than that of divergent.

Mitochondrial functions of de novo proteins would be consistent with much experimental evidence for the cellular roles of microproteins (as most de novo are) and the general preponderance of microprotein association to mitochondria^85–87^. Cases such as BRAWNIN^85^, SMIM4^72^ and MTLN^88^ have established the essential roles of microproteins in mitochondrial respiratory complex assembly and lipid oxidation. Other small mitochondrial proteins function in protein import and translation systems, stress response, apoptosis and more^86^. While most of the above work concerns mammalian cells, small mitochondrial proteins are also known in *S. cerevisiae* and some function in similar pathways^89–92^, yet they are significantly less understood and the connection to de novo gene birth had not been previously made^93^. It is tempting to hypothesize that de novo microproteins originating independently can assume analogous mitochondrial functions in vertebrates and fungi, explaining the absence of homology among them. The crucial insight of our work here is that this seems to apply to most, if not all Saccharomycotina lineages making it a central theme in de novo evolution in yeasts and showing the value of exploratory evolutionary genomics at broad phylogenetic scales.

Our results further suggest that another subset of novel proteins of both origins have endoplasmic reticulum (ER) membrane related functions and in the case of de novo this subset is distinct from the mitochondrial one. Again, many microproteins are known to function in the ER^94^ including in protein translocation and stress response. The ER enrichment, taken together with top molecular function and biological process Gene Ontology terms such as peptide metabolic process and structural constituent of ribosome invokes molecular systems related to the translation of membrane proteins in ER-associated polysomes^95–97^, though these molecular function and biological process terms only concern a small number of genes. Yet another related functional role may be the one found for PIGBOS, a human mitochondrial outer membrane microprotein that interacts at ER-mitochondria contact sites and regulates the unfolded protein response (UPR)^73^. Upregulation of UPR is one hypothesis offered by previous work for the beneficial fitness effects seen when transmembrane de novo proteins are overexpressed in *S. cerevisiae*^28^ and most of these proteins localize to the ER^31^. We hope that our work will serve as a basis for future experimental studies that will validate the expression and function of some of the de novo proteins detected in this study.

Whatever function the de novo proteins we detect here may perform, our analyses suggest that in most lineages they are likely influenced by the repetitive intergenic motifs from which they originate^24,27^. Here we found that polyT motifs are more than three times more common in de novo gene sequences than conserved or divergent genes, and that this is true across genera. Beyond suggesting an intriguing functional and evolutionary interplay of the regulatory role of these motifs and their protein-coding potential (through their translation into TM proteins), our findings demonstrate that such homopolymeric repeats are an important marker when it comes to distinguishing novel gene origination mechanisms.

De novo gene identification is inherently limited by phylogenetic distance and the power of annotation methodologies^98,99^. Both these limitations apply to our study. First, many, perhaps most of de novo genes remain undetected as strong positive evidence for their de novo status is beyond reach due to the absence of close outgroup species. Second, our predictions rely on existing gene annotations, and any annotation methodology ultimately needs to classify an ORF as a true CDS or not. The criteria used, which can change from one version to the next, mean that a sequence can lose its gene status and this can and has been the case for very young, species-specific de novo genes in the past^100^; indeed this is expected as, by definition, they occupy almost the very edge of “gene-ness”. That edge usually comes with neutral or very weak selection^13,101,102^ and is consistent with our own selection analyses (**Figure 2E,G**). The few de novo genes that are under high positive selection comprise a group of genes that might be contributing to rapid adaptation in response to environmental changes^103,104^. Finally, we have done our best to discriminate de novo and divergent origins but the fact that young de novo genes are known to evolve fast suggests that some of the divergent novel genes could be slightly older de novo ones that have rapidly diversified while maintaining synteny.

Structure predictions of de novo proteins also come with important limitations, as others have pointed out^49^. Small proteins (<100 AA in length) pose a difficult challenge when trying to predict their tertiary structure. Generally, protein structures stabilize by their long-range contacts. As smaller molecules inherently have fewer long-range contacts, current state of the art tools, especially single-sequence pLM-based predictors, tend to generate α-helices for such cases, based on sequence propensities^105^. This structure relies on local interactions and also maximizes the pLDDT metric. This leads to high confidence hallucinations that must be dealt with carefully^76^. Lack of sequence similarity in orphan cases means it is generally impossible to evaluate these structures compared to their closest relatives, and they are too distant from the sequence space of the training set. Combined, these challenges mean that structure predictions for small orphan proteins require careful evaluation for reproducibility and the incorporation of more global metrics like pTM alongside pLDDT.

Finally, we believe that our work has implications for our understanding of protein fold evolution and our view of structural similarity. While more work will be necessary, we provide evidence showing that basic protein structural domains can evolve “from scratch” from noncoding DNA. This affords us to raise the question of the deep ancestry all domains belonging to ancient protein folds^41–43,106^, a question which we believe is worth pursuing in the future.

## Materials and Methods

### Data collection

The main Saccharomycotina genomic dataset used was obtained from Shen et al^58^. Within that study, out of a total of 332 genomes, 136 were already publicly available, 24 sequenced by RIKEN and never described before, and 196 were newly sequenced by the Y1000+ Project. Apart from the Saccharomycotina species, additional data from 11 species, 9 from the Pezizomycotina subphylum and 2 from the Taphrinomycotina subphylum, were included and used as outgroups. In addition to these 343 species’ genomes, 10 more species from the division Basidiomycota were randomly selected out of those available in NCBI’s RefSeq in February of 2022 as further outgroups (see Supp. Table 10). The initial dataset of 343 genomes included a collection of predicted coding DNA sequences totaling 2,012,541 sequences, grouped by Shen et al. into orthologous groups (gene/protein families) on the basis of protein sequence similarity using OrthoMCL. We used these families as a starting point for our analyses. The phylogenetic tree that we use is the one produced by Shen et al. by using the non-Bayesian RelTime method (“332_2408OGs_timetree_reltime.nwk” file). The larger genomic dataset of Opulente et al.^64^ consisting of more than 1,000 Saccharomycotina species was also used as a benchmark. The Global Microbial Gene Catalogue (GMGC^107^) v1.0 protein database and the NCBI Bacterial and Archaeal RefSeq database were downloaded in February of 2023.

For intra-specific analyses, we downloaded all available genome assemblies of Saccharomycotina species from NCBI GenBank in January of 2025. Overall, we retrieved 707 assemblies from 137 species. For *S. cerevisiae,* we used 1,086 high quality assemblies^108^.

### Identification of Taxonomically Restricted Genes (TRGs), estimation of timing of origination of gene families and re-clustering

Starting with 233,478 OrthoMCL clusters of Shen et al. we removed those containing any of the following: 1) a member of an outgroup species (Pezizomycotina/Taphrinomycotina), 2) a protein with a statistically significant protein sequence similarity to any protein from 10 randomly sampled Basidiomycota species (DIAMOND BLASTp ultra-sensitive search with default values and E-value cut-off of 0.001), 3) a protein with a statistically significant similarity match to any eukaryotic protein in NCBI’s NR excluding Saccharomycotina (taxid 147537) and Bacteria (taxid 2), using DIAMOND BLASTp ultra-sensitive search with default values and E-value cut-off of 0.001. For each of the remaining 145,417 initial TRG families, we calculated its branch of origin by taking the phylogenetic branch corresponding to the most recent common ancestor (DOLLO parsimony). These families were then re-clustered as follows: we used them as queries to search against the entire protein catalogues with DIAMOND BLASTp in ultra-sensitive mode keeping as significant only alignments with >40% identity and the default e-value (0.001). TRG families were merged, either with other TRG families or with conserved families, if any of their members had a significant match with a member of another family. Note that this re-clustering approach may sometimes lead to the grouping of non-homologous proteins through similarity to different parts of a common protein but we prefer to tolerate more of this error here than its inverse, that is splitting homologous families. Branches of origin were again calculated for merged families and any TRG family merged with a conserved one was then reclassified as conserved.

### Synteny analyses for identification of initial de novo gene candidates and divergent novel genes

To establish syntenic blocks between pairs of Saccharomycotina species we first run MCScanX^109^ with default values for all 54,949 pairwise species combinations. For each initial TRG family we then identified the three closest outgroup clades based on their branch of origin: the first outgroup is the sister clade to the family, the second outgroup is sister clade to the clade containing the family and the first outgroup and similarly for the third outgroup. An outgroup clade can contain one or more species. Based on the MCScanX syntenic blocks we then searched for conserved syntenic regions to every species of each of the three outgroup clades as follows: for each member of the family (referred as “focal gene”), we looked at whether it was found in conserved micro-synteny with respect to each outgroup species by examining the gene order and position of its immediate -1 and +1 neighbors within the corresponding syntenic block identified by MCScanX. Specifically, we established two types of arrangements: (1) the homologues of the -1 and +1 neighbors in the outgroup species are adjacent, found next to each other in the same chromosome and in the same order as the neighbors of the focal gene (homologue of the -1 precedes homologue of the +1), (2) the homologues of the -1 and +1 neighbors are in the same chromosome, in the same order as the neighbors of the focal genes but are separated by one or more genes. A family was considered as an initial de novo candidate, if (a) at least one of its genes was found in a type (1) conserved synteny arrangement with at least one of the species belonging to each one of the three outgroup clades (b) no gene was found in a type (2) conserved synteny arrangement in any of the outgroup species in the three outgroup clades. In total, we were able to identify 6,109 de novo candidate families composed of 6,441 genes. For these families, we extracted for further analysis all intergenic regions of outgroup genomes that are orthologous to family members (the region between the -1 homologue and the +1 homologue for cases found in type (1) conserved synteny arrangement). Based on the same syntenic data we defined novel families that likely resulted through divergence if at least one gene in a family had a type (2) arrangement as previously described with at least one species in at least two out of the three outgroup clades. The entire family was classified as divergent for consistency. For both initial de novo and divergent families only those that did not show additional similarity beyond their ingroup during re-clustering were considered. Finally, we also discarded initial de novo and divergent candidate families containing proteins with any statistically significant similarity hit to any protein from any species outside the family-specific ingroup (DIAMOND BLASTp ultra-sensitive E-value<0.0001). Overall, our criteria ensure that de novo and divergent proteins can be considered equal in terms of sequence novelty.

### Detection of Horizontal Gene Transfers

We conducted three similarity searches using the protein sequences of the initial set of de novo candidates as queries against prokaryotic sequence databases: 1) we searched against the entire GMGC v1.0 protein catalogue clustered at 95% identity (GMGC10.95nr.faa) using DIAMOND BLASTp in very-sensitive mode with options *--max-target-seqs 300* and *--evalue 0.001*, 2) we searched against the bacterial and archaeal NCBI RefSeq protein database using the same parameters as 1, and 3) we searched against “All The Bacteria”^63^ genome database using Lexicmap^110^ with default values and the e-value cut-off set to 0.001. The number of query sequences that mapped against an assembly in ATB was 567. The quality of the assemblies in the database varies, therefore we considered only matches to well-assembled genomes with completeness percentage greater than 90% and contamination less than 5% (Supp. Figure 4). Any query protein with a match in any of the three databases was considered a putative HGT.

In the case of Candida_montana@Seq_485 we used 9 orthologous regions from related species (*B. californica*, *B. hawaiiensis*, *B. populi*, *B. pratensis*, *B. salicaria*, *P. opuntiae*, *P.antillensis*, *P.thermotolerans*, *C.orba*) and 13 more distant species (*K. pastoris*, *W. ciferrii*, *K. populi*, *S. quercuum*, *C. stellimalicola*, *K. ogatae*, *K. pseudopastoris*, *W. bovis*, *W.alni*, *C. azyma*, *W. canadensis*, *S. amethionina*, *W. hampshirensis*) that were selected on the basis of having an ortholog for the upstream and downstream genes of the focal gene. The GC content of the extended region 5kbp up and downstream of the de novo candidate with a sliding window. The step of the sliding window was 100 bp and the length of the window was 1000 bp. To create the trajectories, we kept 100 windows, leaving out 3 or 4 per case, so that all genes have the same number and then we averaged the GC per window per gene. One gene, that belonged to the closely related category, was left out because it was short, it had only 87 windows.

### Estimation of false positives in initial de novo gene candidate set

To assess the false positive proportion of our initial de novo candidate dataset we used a larger Saccharomycotina dataset^64^, that was comprised of 1,154 genomes. The de novo candidates were used as queries for a protein similarity search using DIAMOND against the proteome of the whole dataset, which consisted of 6,830,348 proteins, using the ultra-sensitive mode. If homology was found in a species X outside of its sister species, and the last common ancestor with that species X was older than the age which the candidate was assigned, then its age was changed to the same as the last common ancestor with species X. The de novo candidate protein sequences were, also, mapped against the genomes, using miniprot^111^ with the default settings. For these mappings we set some thresholds in order to consider them as real homology: identity over 70%, coverage over 40%. As closely related non-coding sequences can easily exceed these criteria, we only considered cases where the last common ancestor of the species that the protein mapped on should be older than the last common ancestor with the furthest outgroup species used to qualify a gene as candidate de novo (see **Figure 1**).

### Gene and protein properties

CDS lengths were calculated with the *faSize* utility downloaded from http://hgdownload.soe.ucsc.edu/admin/exe/ using the CDS FASTA files provided by Shen et al. GC content and GRAVY scale hydropathicity values were calculated using the program CodonW. To acquire predictions about the nucleosomal positioning on the Saccharomycotina genomes, the algorithm SymCurv^112^ was used. SymCurv is a computational *ab initio* method for nucleosome positioning prediction. It is based on the structural property of natural nucleosome forming sequences, to be symmetrically curved around a local minimum of curvature. The FASTA files of the whole genomes were used as an input and the output that was kept were the final predictions. The nucleotide coverage was calculated by using the gene coordinates found in the GTF files which were intersected with the nucleotide positioning coordinates, which were predicted as described above, with the use of intersect of the BED suite tools^113^. The SeqUtils.IsoelectricPoint python package was used for the calculation of the isoelectric point of the proteins, from the Bio module. The biosynthetic cost of each protein was calculated using a custom Python script with the use of a cost per amino acid matrix that was published by Raiford et al^114^. Phobius^115^ was used to predict transmembrane domains and PredictProperty^116^ was used to predict secondary structure elements (ss3 output) and solvent accessibility. The tool IUPRED3^117^ with the *-long* parameter was used to predict intrinsically disordered regions on protein sequences. DeepLoc 1^118^ was used to predict subcellular localization.

Subcellular localization was predicted using DeepLoc (version 1)^71^. We created the two control sets with custom python scripts. One randomly shuffled the de novo set, and the other cut random segments of randomly chosen conserved proteins matching the exact same length of the de novo protein set.

### Calculation of signatures of selection

To estimate selection pressure from intra-specific alignments we first identified the locus of every de novo gene’s CDS in every genome assembly of the same species, downloaded from NCBI. Regions corresponding to the CDS were identified using BLASTn and then extracted and then a multiple sequence alignment was generated (including the query CDS) using MAFFT in *– auto* mode. The alignments were then filtered using GoAlign^119^ to remove duplicate sequences and alignments with more than 30% gaps on average. Then, using the trim tool of the GoAlign suite, alignments with a sequence length that was not perfectly divided by three were trimmed so as to have a number of bases divided perfectly by three. The RaxML-ng^120^ tool was run on the alignment files having more than four alignments, using the following command:

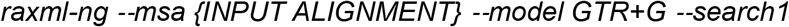

The trees produced by RaxML were used to run HyPhy on to calculate single alignment omega values (*d_N_/d_S_*) by running the following command:

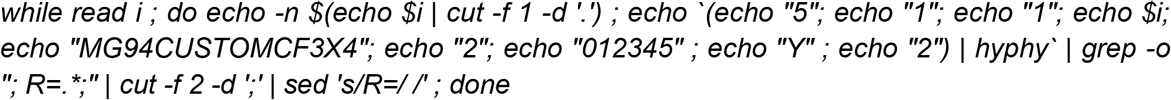

The above command outputs the omega values for each of the alignment files given as input. As a positive control, we used short conserved sequences: we picked out the genes of 30 conserved genes’ families with the shortest mean length. The mean sequence length value of the selected conserved genes’ CDS was 593.2 nt. Intraspecific alignments were then generated in the same manner as for de novo genes and the same filtering and commands were run to calculate the omega values. To produce the set of shuffled sequences, the *multiperm*^121^ tool was run on the denovo intraspecific alignments as follows: *multiperm --conservation=none -w {Input Alignment File}*. In-frame stop codons where then removed from the shuffled alignments using a custom Python script and RaxML and HyPhy tools were run the same way as for the de novo dataset.

For selection signatures on inter-specific alignments we performed the following: multiple sequence alignments were generated using MAFFT --auto for every multispecies de novo gene family and then these alignments were used as input to PAML^122^ *yn00*. We used the Yang & Nielsen (2000) method and kept only pairwise comparisons where *d_S_* > 0 and standard error of *d_S_* is less than half of the value of *d_S_*. One value per family was then obtained by calculating the average *d_N_/d_S_* (omega) out of all remaining pairwise comparisons.

### Motif searches

We used FIMO of the MEME^123^ suite to search for occurrences of the three motifs identified by Vakirlis and Fuqua^29^ within entire Saccharomycotina genome sequences and separately within CDS sequences:

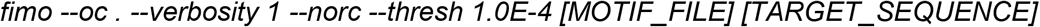

Note that using the *–norc* option only the forward strand is searched.

To calculate the number of bp in coding and non-coding regions that overlap with the predicted motifs we used *intersect* from the bedtools suite^113^ and the GTF file of each species. The regions that overlapped multiple times with the motifs were filtered and counted only once. To calculate the motif density difference between the three different motifs we used the following formula:

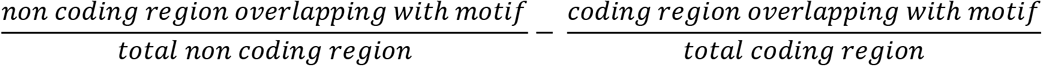

To calculate the motif occurrence of each motif in the different types of genes (de novo, divergent, TRG and conserved) we summed the total number of nucleotides overlapping with each motif for every gene type and the normalized with the sum of the length of all the genes of each type. So for each motif per gene type we followed this formula:

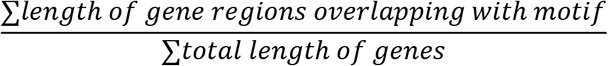

### Tertiary protein structure prediction

Models of tertiary structures of de novo and divergent proteins were generated using ESM3 3.2.1.post1^74^ on a NVIDIA GeForce RTX 5090. As ESM3 is a generative model, we decided to run 10 replicates per protein sequence, with the model esm3-sm-open-v1^124^. Seed was set to 42, number of steps to 32, temperature set to default value (=1) and besides the .PDB output files we retained average pLDDT and pTM scores per generated structure. As a control, we shuffled each de novo and divergent protein sequence with a custom python script and generated structures in the same way as for the original sequences. Using seqkit 2.13^125^, we generated 5,185 random sequences with the same length distribution as that of de novo proteins and generated structures for those as well. Because structural hallucinations can have high confidence metrics and small proteins are not always bound to a dominant fold but can have a range of microstates, before we filtered the generated structures, we clustered the 10 generated structures per protein with Foldseek 10.941cd33^126^ easy-search. TM threshold was 0.5 and coverage 0.6. If a protein had a cluster with a size of at least 5 members (50%), meaning at least 5 of the 10 replicates produce the same structure, we assume that this generated structure is reproducible by esm3. After the clustering step and the selection of the main cluster per protein, we calculate the average pLDDT x pTM only for the members of that cluster and retain the single structure with the highest value as the best structural representative of the cluster. These remaining structures are then filtered for pTM >=0.8 and pLDDT>=0.7 and are considered High Confidence (HC) structures. By talking advantage of the generative nature of the model, we can with these steps isolate proteins showing a high confidence dominant fold, from others who either jump between microstates and a single-fold prediction would be considered an underrepresentation of its conformation space, or from those whose sequence lead to hallucinations and would otherwise pass filtering. Lastly, we generated Secondary Structure predictions for each HC structure using their PDB files as input for ESM3. Protein structures were visualized using py3Dmol 2.5.3^78^.

### Structural similarity searches and associated analyses of entire proteins and domains

Using Foldseek easy-search we compared our high confidence (HC) dataset against afdb50 v4^127^ and mgnify-esm30 v0^128^. TM cutoff was set at 0.4, e-value at 0.001, probability >=0.5 and sensitivity = 9,5. Cov-mode was set to 5. Cases of interest with a match were then re-run this time on the Foldseek server, isolating the superpositioned structures of the queries and their best hit along with the alignment TM score and RMSD. We segmented the high confidence folds into domains and compared them against TED v5^77^ with the use of merizo-search^82^ easy-search. Cov-mode was set to 2, coverage at 0.6 and TM threshold of 0.5. Predicted domains were then filtered for pIoU >=0.75 and hits for prob>=0.5. The target domains of the remaining hits were used to map 285/369 queries to 75 unique CATH labels. The queries with no match in the Foldseek and the merizo-search analysis are considered novel folds.

For each HC structure of de novo and divergent novel proteins we predicted GO terms using their PDB files as input for DeepFRI 1.0.0^21^. We also predicted GO terms for the complete list of de novo and divergent genes, using their amino acid sequences this time as input. The two approaches produced highly similar results.

To detect similarity of divergent novel proteins with HC structures to their “invisible” homologues we performed the following: we generated the structures of the “invisible” homologues as described above and only HC structures were kept using the exact same criteria. Foldseek easy-search was used to compare the structures of all the divergent HC structures against all the respective homologues. E-value was set to 0.001, cov-mode to 5, TM threshold at 0.3 and sensitivity at 9.5. The output was filtered for prob>=0.5.

Using esm3 cookbook for UMAPs^130^, we generated embeddings for 286,243 conserved proteins, 278 most ancient TRGs, 5,126 species-specific de novo and 4,394 species-specific divergent novel proteins. We projected these into a two-dimensional space, using Python’s umap 0.5.12^131^.

### Quantification and statistical analysis

All statistics were done in R version 4.3.3. Plots were generated using ggplot2^132^. All statistical details including the type of statistical test performed and exact value of n can be found in the Results and figure legends. Boxplots show median (horizontal line inside the box), first and third quartiles of data (lower and upper hinges) and values no further or lower than 1.5*distance between the first and third quartiles (upper and lower whisker). No methods were used to determine a priori whether the data met assumptions of the statistical approaches. Alignments were visualized with the Jalview software^133^.

## Data availability

All analyses performed used publicly available data. All data produced by analyses of this study are available in the source data of this article and/or in this GitHub [repository](https://github.com/emiliostassios/Evolutionary-origins-of-molecular-novelty-across-an-entire-yeast-subphylum-).

## Acknowledgements

We are deeply grateful to Dr. Timothy Fuqua for his critical reading and feedback on this manuscript. The research project was supported by the Hellenic Foundation for Research and Innovation (H.F.R.I.) under the “3rd Call for H.F.R.I. Research Projects to support Post-Doctoral Researchers” to N. Vakirlis (Project Number:7330) and by a G4 grant from Institut Pasteur awarded to N. Vakirlis. This work was supported by a PhD scholarship by a Fondation Santé “Sidney Altman Scholarship Program” to E. Tassios. E. Tassios was partially supported by a BSRC “Alexander Fleming” Installation Grant to C. Nikolaou (Programme COMPETIΤΙVENESS, NSRF 2021-2027). This project was partially supported by the National Science Foundation under grants nos. [DEB-2110403](https://www.sciencedirect.com/unsupported_browser#gs1) (to C. T. Hittinger) and [DEB-2110404](https://www.sciencedirect.com/unsupported_browser#gs1) (to A. Rokas); the Great Lakes Bioenergy Research Center, U.S. Department of Energy, Office of Science, Biological and Environmental Research Program under Award Number DE-SC0018409 (of which C. T. Hittinger is a co-investigator); and the National Institute of Food and Agriculture, United States Department of Agriculture, Hatch project 7005101 (to C. T. Hittinger); and the Vilas Trust Estate (to C. T. Hittinger).

## Author information

### Contributions

NV conceived the study. ET, NV, EMT and DR performed the analyses and analysed the data. NV, CN, AR and CTH supervised the work. NV and CN acquired funding for the work. NV and ET wrote the initial draft. NV, CN, AR and ET revised the initial draft. All authors read, finalized and approved the final manuscript.

### Corresponding authors

Correspondence to Nikolaos Vakirlis and Christoforos Nikolaou

## References

1. Dujon, B. The yeast genome project: what did we learn? Trends Genet. TIG 12, 263–270 (1996).

2. Vakirlis, N., Carvunis, A.-R. & McLysaght, A. Synteny-based analyses indicate that sequence divergence is not the main source of orphan genes. eLife 9, e53500 (2020).

3. Prabh, N. & Tautz, D. Frequent lineage-specific substitution rate changes support an episodic model for protein evolution. *G3* GenesGenomesGenetics 11, jkab333 (2021).

4. Wolfe, K. Evolutionary Genomics: Yeasts Accelerate beyond BLAST. Curr. Biol. 14, R392–R394 (2004).

5. Andersson, D. I., Jerlström-Hultqvist, J. & Näsvall, J. Evolution of New Functions De Novo and from Preexisting Genes. Cold Spring Harb. Perspect. Biol. 7, a017996 (2015).

6. Tassios, E., de Leuw, J., Nikolaou, C., Kupczok, A. & Vakirlis, N. Machine learning can distinguish orphans that have resulted from sequence divergence beyond recognition. Bioinforma. Adv. vbaf324 (2025) doi:10.1093/bioadv/vbaf324.

7. Weisman, C. M., Murray, A. W. & Eddy, S. R. Many, but not all, lineage-specific genes can be explained by homology detection failure. PLOS Biol. 18, e3000862 (2020).

8. Prabh, N. & Rödelsperger, C. De Novo, Divergence, and Mixed Origin Contribute to the Emergence of Orphan Genes in Pristionchus Nematodes. G3 GenesGenomesGenetics 9, 2277–2286 (2019).

9. Pereira, A. B. et al. Orphan genes are not a distinct biological entity. Bioessays 47, 2400146 (2025).

10. Tautz, D. & Domazet-Lošo, T. The evolutionary origin of orphan genes. Nat. Rev. Genet. 12, 692– 702 (2011).

11. Arendsee, Z. W., Li, L. & Wurtele, E. S. Coming of age: orphan genes in plants. Trends Plant Sci. 19, 698–708 (2014).

12. Colson, P., La Scola, B., Levasseur, A., Caetano-Anollés, G. & Raoult, D. Mimivirus: leading the way in the discovery of giant viruses of amoebae. Nat. Rev. Microbiol. 15, 243–254 (2017).

13. Boratto, P. V. M. et al. Yaravirus: A novel 80-nm virus infecting Acanthamoeba castellanii. Proc. Natl. Acad. Sci. U. S. A. 117, 16579–16586 (2020).

14. Van Oss, S. B. & Carvunis, A.-R. De novo gene birth. PLOS Genet. 15, e1008160 (2019).

15. Baalsrud, H. T. et al. De Novo Gene Evolution of Antifreeze Glycoproteins in Codfishes Revealed by Whole Genome Sequence Data. Mol. Biol. Evol. 35, 593–606 (2018).

16. Li, L. et al. Identification of the novel protein QQS as a component of the starch metabolic network in Arabidopsis leaves. Plant J. 58, 485–498 (2009).

17. Xiao, W. et al. A Rice Gene of De Novo Origin Negatively Regulates Pathogen-Induced Defense Response. PLoS ONE 4, e4603 (2009).

18. Ritschard, E. A. et al. Coupled Genomic Evolutionary Histories as Signatures of Organismal Innovations in Cephalopods. BioEssays 0, 1900073.

19. Blevins, W. R. et al. Uncovering de novo gene birth in yeast using deep transcriptomics. Nat. Commun. 12, 604 (2021).

20. Vakirlis, N. et al. A Molecular Portrait of De Novo Genes in Yeasts. Mol. Biol. Evol. 35, 631–645 (2018).

21. Prabh, N. et al. Deep taxon sampling reveals the evolutionary dynamics of novel gene families in Pristionchus nematodes. Genome Res. 28, 1664–1674 (2018).

22. Zhang, L. et al. Rapid evolution of protein diversity by de novo origination in Oryza. *Nat*. Ecol. Evol. 3, 679–690 (2019).

23. Ruiz-Orera, J., Verdaguer-Grau, P., Villanueva-Cañas, J. L., Messeguer, X. & Albà, M. M. Translation of neutrally evolving peptides provides a basis for de novo gene evolution. Nat. Ecol. Evol. 1 (2018) doi:10.1038/s41559-018-0506-6.

24. Carvunis, A.-R. et al. Proto-genes and de novo gene birth. Nature 487, 370–374 (2012).

25. Cai, J., Zhao, R., Jiang, H. & Wang, W. *De Novo* Origination of a New Protein-Coding Gene in *Saccharomyces cerevisiae*. Genetics 179, 487–496 (2008).

26. Wacholder, A. et al. A vast evolutionarily transient translatome contributes to phenotype and fitness. Cell Syst. 14, 363–381.e8 (2023).

27. Montañés, J. C., Huertas, M., Messeguer, X. & Albà, M. M. Evolutionary Trajectories of New Duplicated and Putative De Novo Genes. Mol. Biol. Evol. 40, msad098 (2023).

28. Tassios, E., Nikolaou, C. & Vakirlis, N. Intergenic Regions of Saccharomycotina Yeasts are Enriched in Potential to Encode Transmembrane Domains. Mol. Biol. Evol. 40, msad059 (2023).

29. Papadopoulos, C. et al. Intergenic ORFs as elementary structural modules of de novo gene birth and protein evolution. Genome Res. 31, 2303–2315 (2021).

30. Roginski, P. et al. Impact of GC content on de novo gene birth. Nat. Commun. 17, 1268 (2026).

31. Vakirlis, N. et al. De novo emergence of adaptive membrane proteins from thymine-rich genomic sequences. Nat. Commun. 11, 781 (2020).

32. Vakirlis, N. & Fuqua, T. Intergenic polyA/T tracts explain the propensity of yeast de novo genes to encode transmembrane domains. J. Evol. Biol. voaf089 (2025) doi:10.1093/jeb/voaf089.

33. Tsankov, A. M., Thompson, D. A., Socha, A., Regev, A. & Rando, O. J. The Role of Nucleosome Positioning in the Evolution of Gene Regulation. PLOS Biol. 8, e1000414 (2010).

34. Fuqua, T. & Vakirlis, N. Emergence Biases in Molecular Evolution. Genome Biol. Evol. 18, evag195 (2026).

35. Li, D. et al. A de novo originated gene depresses budding yeast mating pathway and is repressed by the protein encoded by its antisense strand. Cell Res. 20, 408–420 (2010).

36. Li, D., Yan, Z., Lu, L., Jiang, H. & Wang, W. Pleiotropy of the de novo-originated gene MDF1. Sci. Rep. 4, 7280 (2014).

37. Houghton, C. J. et al. Cellular processing of beneficial de novo emerging proteins. bioRxiv 2024.08.28.610198 (2024) doi:10.1101/2024.08.28.610198.

38. Zhao, L., Svetec, N. & Begun, D. J. De Novo Genes. Annu. Rev. Genet. 58, 211–232 (2024).

39. Bornberg-Bauer, E. & Eicholt, L. A. Emergence and evolution of protein-coding de novo genes. Nat. Rev. Genet. 27, 530–546 (2026).

40. Gligorijević, V. et al. Structure-based protein function prediction using graph convolutional networks. Nat. Commun. 12, 3168 (2021).

41. Sarkar, A., Krishnan, K. & Eddy, S. R. Protein Sequence Domain Annotation using Language Models. 2024.06.04.596712 Preprint at 10.1101/2024.06.04.596712 (2024).

42. Littmann, M., Heinzinger, M., Dallago, C., Olenyi, T. & Rost, B. Embeddings from deep learning transfer GO annotations beyond homology. Sci. Rep. 11, 1160 (2021).

43. Barrios-Núñez, I., et al. Decoding functional proteome information in model organisms using protein language models. NAR Genomics Bioinforma. 6, lqae078 (2024).

44. Jumper, J. et al. Highly accurate protein structure prediction with AlphaFold. Nature 596, 583–589 (2021).

45. Lin, Z. et al. Evolutionary-scale prediction of atomic-level protein structure with a language model. Science 379, 1123–1130 (2023).

46. Hochberg, G. K. A. & Thornton, J. W. Reconstructing Ancient Proteins to Understand the Causes of Structure and Function. Annu. Rev. Biophys. 46, 247–269 (2017).

47. Koonin, E. V., Wolf, Y. I. & Karev, G. P. The structure of the protein universe and genome evolution. Nature 420, 218–223 (2002).

48. Alvarez-Carreño, C. Piecing Together the History of Protein Folds From a Fragmented Evolutionary Record. Genome Biol. Evol. 17, evaf148 (2025).

49. Alva, V., Söding, J. & Lupas, A. N. A vocabulary of ancient peptides at the origin of folded proteins. eLife 4, e09410 (2015).

50. Lupas, A. N., Ponting, C. P. & Russell, R. B. On the Evolution of Protein Folds: Are Similar Motifs in Different Protein Folds the Result of Convergence, Insertion, or Relics of an Ancient Peptide World? J. Struct. Biol. 134, 191–203 (2001).

51. Bornberg-Bauer, E., Hlouchova, K. & Lange, A. Structure and function of naturally evolved de novo proteins. Curr. Opin. Struct. Biol. 68, 175–183 (2021).

52. Lange, A. et al. Structural and functional characterization of a putative de novo gene in Drosophila. Nat. Commun. 12, 1667 (2021).

53. Middendorf, L., Ravi Iyengar, B. & Eicholt, L. A. Sequence, Structure, and Functional Space of Drosophila De Novo Proteins. Genome Biol. Evol. 16, evae176 (2024).

54. Peng, J. & Zhao, L. The origin and structural evolution of de novo genes in Drosophila. Nat. Commun. 15, 810 (2024).

55. Aubel, M., Eicholt, L. & Bornberg-Bauer, E. Assessing structure and disorder prediction tools for de novo emerged proteins in the age of machine learning. F1000Research 12, 347 (2023).

56. Seçkin, E., Colinet, D., Danchin, E. G. J. & Sarti, E. Evaluating transformer-based models for structural characterization of orphan proteins. Bioinforma. Adv. vbag215 (2026) doi:10.1093/bioadv/vbag215.

57. Liu, J. et al. Do “Newly Born” orphan proteins resemble “Never Born” proteins? A study using three deep learning algorithms. Proteins Struct. Funct. Bioinforma. 91, 1097–1115 (2023).

58. Durairaj, J. et al. Uncovering new families and folds in the natural protein universe. Nature 622, 646– 653 (2023).

59. Koehler Leman, J., et al. Sequence-structure-function relationships in the microbial protein universe. Nat. Commun. 14, 2351 (2023).

60. Nomburg, J. et al. Birth of protein folds and functions in the virome. Nature 633, 710–717 (2024).

61. Allen, J. O. et al. Comparisons among two fertile and three male-sterile mitochondrial genomes of maize. Genetics 177, 1173–1192 (2007).

62. Zhuang, X., Yang, C., Murphy, K. R. & Cheng, C.-H. C. Molecular mechanism and history of non-sense to sense evolution of antifreeze glycoprotein gene in northern gadids. Proc. Natl. Acad. Sci. U. S. A. 116, 4400–4405 (2019).

63. Shen, X.-X. et al. Tempo and Mode of Genome Evolution in the Budding Yeast Subphylum. Cell 175, 1533–1545.e20 (2018).

64. Basile, W., Sachenkova, O., Light, S. & Elofsson, A. High GC content causes orphan proteins to be intrinsically disordered. PLOS Comput. Biol. 13, e1005375 (2017).

65. Vakirlis, N. & McLysaght, A. Computational Prediction of De Novo Emerged Protein-Coding Genes. in Computational Methods in Protein Evolution (ed. Sikosek, T.) vol. 1851 63–81 (Springer New York, New York, NY, 2019).

66. Gonçalves, C., Hittinger, C. T. & Rokas, A. Horizontal Gene Transfer in Fungi and Its Ecological Importance. in Fungal Associations (eds Hsueh, Y.-P. & Blackwell, M.) 59–81 (Springer International Publishing, Cham, 2024). doi:10.1007/978-3-031-41648-4_3.

67. Hunt, M., Lima, L., Shen, W., Lees, J. & Iqbal, Z. AllTheBacteria - all bacterial genomes assembled, available and searchable. 2024.03.08.584059 Preprint at 10.1101/2024.03.08.584059 (2024).

68. Opulente, D. A. et al. Genomic factors shape carbon and nitrogen metabolic niche breadth across Saccharomycotina yeasts. Science 384, eadj4503 (2024).

69. Vakirlis, N. & Kupczok, A. Large-scale investigation of species-specific orphan genes in the human gut microbiome elucidates their evolutionary origins. Genome Res. 34, 888–903 (2024).

70. Kryazhimskiy, S. & Plotkin, J. B. The Population Genetics of dN/dS. PLOS Genet. 4, e1000304 (2008).

71. Xia, S., Chen, J., Arsala, D., Emerson, J. J. & Long, M. Functional innovation through new genes as a general evolutionary process. Nat. Genet. 57, 295–309 (2025).

72. Chen, S., Zhang, Y. E. & Long, M. New Genes in Drosophila Quickly Become Essential. Science 330, 1682–1685 (2010).

73. Echols, N. et al. Comprehensive analysis of amino acid and nucleotide composition in eukaryotic genomes, comparing genes and pseudogenes. Nucleic Acids Res. 30, 2515–2523 (2002).

74. Wissler, L., Gadau, J., Simola, D. F., Helmkampf, M. & Bornberg-Bauer, E. Mechanisms and Dynamics of Orphan Gene Emergence in Insect Genomes. Genome Biol. Evol. 5, 439–455 (2013).

75. Liang, C. et al. Mitochondrial microproteins link metabolic cues to respiratory chain biogenesis. Cell Rep. 40, (2022).

76. Chu, Q. et al. Regulation of the ER stress response by a mitochondrial microprotein. Nat. Commun. 10, 4883 (2019).

77. Brown, C. J., Johnson, A. K., Dunker, A. K. & Daughdrill, G. W. Evolution and Disorder. Curr. Opin. Struct. Biol. 21, 441–446 (2011).

78. Brown, C. J., Johnson, A. K. & Daughdrill, G. W. Comparing Models of Evolution for Ordered and Disordered Proteins. Mol. Biol. Evol. 27, 609–621 (2010).

79. Simulating 500 million years of evolution with a language model | Science. https://www.science.org/doi/10.1126/science.ads0018.

80. Exploring structural diversity across the protein universe with The Encyclopedia of Domains | Science. https://www.science.org/doi/10.1126/science.adq4946.

81. van Kempen, M. et al. Fast and accurate protein structure search with Foldseek. Nat. Biotechnol. 42, 243–246 (2024).

82. Kajava, A. V. & Steven, A. C. Beta-rolls, beta-helices, and other beta-solenoid proteins. Adv. Protein Chem. 73, 55–96 (2006).

83. Mesdaghi, S., Price, R. M., Madine, J. & Rigden, D. J. Deep Learning-based structure modelling illuminates structure and function in uncharted regions of β-solenoid fold space. J. Struct. Biol. 215, 108010 (2023).

84. Cellular processing of beneficial de novo emerging proteins | bioRxiv. https://www.biorxiv.org/content/10.1101/2024.08.28.610198v1.

85. Wang, K. et al. Structural basis for antibiotic resistance by chloramphenicol acetyltransferase type A in Staphylococcus aureus. Sci. Rep. 15, 37020 (2025).

86. Arakawa, T. & Timasheff, S. N. [3]Theory of protein solubility. in Methods in Enzymology vol. 114 49–77 (Elsevier, 1985).

87. Tokmakov, A. A., Kurotani, A. & Sato, K.-I. Protein pI and Intracellular Localization. Front. Mol. Biosci. 8, 775736 (2021).

88. Valli, M. et al. Intracellular pH Distribution in *Saccharomyces cerevisiae* Cell Populations, Analyzed by Flow Cytometry. Appl. Environ. Microbiol. 71, 1515–1521 (2005).

89. Zhang, S. et al. Mitochondrial peptide BRAWNIN is essential for vertebrate respiratory complex III assembly. Nat. Commun. 11, 1312 (2020).

90. Kamradt, M. L. & Makarewich, C. A. Mitochondrial microproteins: critical regulators of protein import, energy production, stress response pathways, and programmed cell death. Am. J. Physiol. - Cell Physiol. 325, C807–C816 (2023).

91. Borghol, N., Yandiev, S. & Courchet, J. Mitochondrial Microproteins: Emerging Regulators in Neurodevelopment and Neurodegeneration. Bioessays 47, e70058 (2025).

92. Stein, C. S. et al. Mitoregulin: A lncRNA-Encoded Microprotein that Supports Mitochondrial Supercomplexes and Respiratory Efficiency. Cell Rep. 23, 3710–3720.e8 (2018).

93. Kastenmayer, J. P. et al. Functional genomics of genes with small open reading frames (sORFs) in S. cerevisiae. Genome Res. 16, 365–373 (2006).

94. Strecker, V. et al. Supercomplex-associated Cox26 protein binds to cytochrome c oxidase. Biochim. Biophys. Acta 1863, 1643–1652 (2016).

95. Busto, J. V. et al. Role of the small protein Mco6 in the mitochondrial sorting and assembly machinery. 2023.02.03.527057 Preprint at 10.1101/2023.02.03.527057 (2023).

96. Esser, K., Pratje, E. & Michaelis, G. SOM 1, a small new gene required for mitochondrial inner membrane peptidase function in Saccharomyces cerevisiae. Mol. Gen. Genet. MGG 252, 437–445 (1996).

97. Parikh, S. B., Houghton, C., Van Oss, S. B., Wacholder, A. & Carvunis, A.-R. Origins, evolution, and physiological implications of de novo genes in yeast. Yeast 39, 471–481 (2022).

98. Coughlin, T. M. & Makarewich, C. A. Emerging Roles for Microproteins as Critical Regulators of Endoplasmic Reticulum Function and Cellular Homeostasis. Semin. Cell Dev. Biol. 170, 103608 (2025).

99. Frey, S., Pool, M. & Seedorf, M. Scp160p, an RNA-binding, polysome-associated protein, localizes to the endoplasmic reticulum of Saccharomyces cerevisiae in a microtubule-dependent manner. J. Biol. Chem. 276, 15905–15912 (2001).

100. Jüschke, C., Ferring, D., Jansen, R.-P. & Seedorf, M. A novel transport pathway for a yeast plasma membrane protein encoded by a localized mRNA. Curr. Biol. CB 14, 406–411 (2004).

101. Baum, S., Bittins, M., Frey, S. & Seedorf, M. Asc1p, a WD40-domain containing adaptor protein, is required for the interaction of the RNA-binding protein Scp160p with polysomes. Biochem. J. 380, 823–830 (2004).

102. Weisman, C. M., Murray, A. W. & Eddy, S. R. Mixing genome annotation methods in a comparative analysis inflates the apparent number of lineage-specific genes. Curr. Biol. CB 32, 2632–2639.e2 (2022).

103. Barrera-Redondo, J., Lotharukpong, J. S., Drost, H.-G. & Coelho, S. M. Uncovering gene-family founder events during major evolutionary transitions in animals, plants and fungi using GenEra. Genome Biol. 24, 54 (2023).

104. McLysaght, A. & Hurst, L. D. Open questions in the study of de novo genes: what, how and why. Nat. Rev. Genet. 17, 567–578 (2016).

105. Ruiz-Orera, J., Verdaguer-Grau, P., Villanueva-Cañas, J. L., Messeguer, X. & Albà, M. M. Translation of neutrally evolving peptides provides a basis for de novo gene evolution. Nat. Ecol. Evol. 2, 890– 896 (2018).

106. Durand, É. et al. Turnover of ribosome-associated transcripts from de novo ORFs produces gene-like characteristics available for de novo gene emergence in wild yeast populations. Genome Res. 29, 932– 943 (2019).

107. Liu, X. et al. Detecting and characterizing genomic signatures of positive selection in global populations. Am. J. Hum. Genet. 92, 866–881 (2013).

108. Luisi, P. et al. Recent Positive Selection Has Acted on Genes Encoding Proteins with More Interactions within the Whole Human Interactome. Genome Biol. Evol. 7, 1141–1154 (2015).

109. A Comprehensive Evaluation of Protein Structure Prediction Models for Short Peptides. Sciety https://sciety.org/articles/activity/10.64898/2026.07.02.736085 (2026).

110. Caetano-Anollés, K., Aziz, M. F., Mughal, F. & Caetano-Anollés, G. On Protein Loops, Prior Molecular States and Common Ancestors of Life. J. Mol. Evol. 92, 624–646 (2024).

111. Coelho, L. P. et al. Towards the biogeography of prokaryotic genes. Nature 601, 252–256 (2022).

112. Loegler, V. et al. From genotype to phenotype with 1,086 near telomere-to-telomere yeast genomes. Nature 1–10 (2025) doi:10.1038/s41586-025-09637-0.

113. Wang, Y. et al. MCScanX: a toolkit for detection and evolutionary analysis of gene synteny and collinearity. Nucleic Acids Res. 40, e49–e49 (2012).

114. Shen, W., Lees, J. A. & Iqbal, Z. Efficient sequence alignment against millions of prokaryotic genomes with LexicMap. Nat. Biotechnol. 1–8 (2025) doi:10.1038/s41587-025-02812-8.

115. Li, H. Protein-to-genome alignment with miniprot. Bioinformatics 39, btad014 (2023).

116. Nikolaou, C., Althammer, S., Beato, M. & Guigó, R. Structural constraints revealed in consistent nucleosome positions in the genome of S. cerevisiae. Epigenetics Chromatin 3, 20 (2010).

117. Quinlan, A. R. & Hall, I. M. BEDTools: a flexible suite of utilities for comparing genomic features. Bioinforma. Oxf. Engl. 26, 841–842 (2010).

118. Raiford, D. W. et al. Do amino acid biosynthetic costs constrain protein evolution in Saccharomyces cerevisiae? J. Mol. Evol. 67, 621–630 (2008).

119. Käll, L., Krogh, A. & Sonnhammer, E. L. L. Advantages of combined transmembrane topology and signal peptide prediction--the Phobius web server. Nucleic Acids Res. 35, W429–432 (2007).

120. Peng, J. & Xu, J. RaptorX: exploiting structure information for protein alignment by statistical inference. Proteins 79, 161–171 (2011).

121. Erdős, G., Pajkos, M. & Dosztányi, Z. IUPred3: prediction of protein disorder enhanced with unambiguous experimental annotation and visualization of evolutionary conservation. Nucleic Acids Res. 49, W297–W303 (2021).

122. Almagro Armenteros, J. J., Sønderby, C. K., Sønderby, S. K., Nielsen, H. & Winther, O. DeepLoc: prediction of protein subcellular localization using deep learning. Bioinformatics 33, 3387–3395 (2017).

123. Lemoine, F. & Gascuel, O. Gotree/Goalign: toolkit and Go API to facilitate the development of phylogenetic workflows. NAR Genomics Bioinforma. 3, (2021).

124. Kozlov, A. M., Darriba, D., Flouri, T., Morel, B. & Stamatakis, A. RAxML-NG: a fast, scalable and user-friendly tool for maximum likelihood phylogenetic inference. Bioinformatics 35, 4453–4455 (2019).

125. Anandam, P., Torarinsson, E. & Ruzzo, W. L. Multiperm: shuffling multiple sequence alignments while approximately preserving dinucleotide frequencies. Bioinformatics 25, 668–669 (2009).

126. Yang, Z. PAML 4: Phylogenetic Analysis by Maximum Likelihood. Mol. Biol. Evol. 24, 1586–1591 (2007).

127. Grant, C. E., Bailey, T. L. & Noble, W. S. FIMO: scanning for occurrences of a given motif. Bioinformatics 27, 1017–1018 (2011).

128. ESM3 Model Family - a EvolutionaryScale Collection. https://huggingface.co/collections/EvolutionaryScale/esm3-model-family (2026).

129. Shen, W., Sipos, B. & Zhao, L. SeqKit2: A Swiss army knife for sequence and alignment processing. iMeta 3, e191 (2024).

130. van Kempen, M. et al. Fast and accurate protein structure search with Foldseek. Nat. Biotechnol. 42, 243–246 (2024).

131. Barrio-Hernandez, I. et al. Clustering predicted structures at the scale of the known protein universe. Nature 622, 637–645 (2023).

132. Lin, Z., et al. ESM Atlas v0 representative random sample of predicted protein structures. OpenAIRE - Explore https://explore.openaire.eu/search/dataset?pid=10.5281%2Fzenodo.7623482 (2022).

133. Kandathil, S. M., Lau, A. M., Buchan, D. W. A. & Jones, D. T. Foldclass and Merizo-search: scalable structural similarity search for single-and multi-domain proteins using geometric learning. Bioinformatics 41, btaf277 (2025).

134. esm/cookbook/tutorials at main · Biohub/esm. GitHub https://github.com/Biohub/esm/tree/main/cookbook/tutorials.

135. McInnes, L., Healy, J. & Melville, J. UMAP: Uniform Manifold Approximation and Projection for Dimension Reduction. Preprint at 10.48550/arXiv.1802.03426 (2020).

136. Wickham, H. ggplot2. WIREs Comput. Stat. 3, 180–185 (2011).

137. Troshin, P. V., Procter, J. B. & Barton, G. J. Java bioinformatics analysis web services for multiple sequence alignment—JABAWS:MSA. Bioinformatics 27, 2001–2002 (2011).

