## Supplementary Figures for "Evolutionary origins of protein novelty across an entire yeast subphylum"

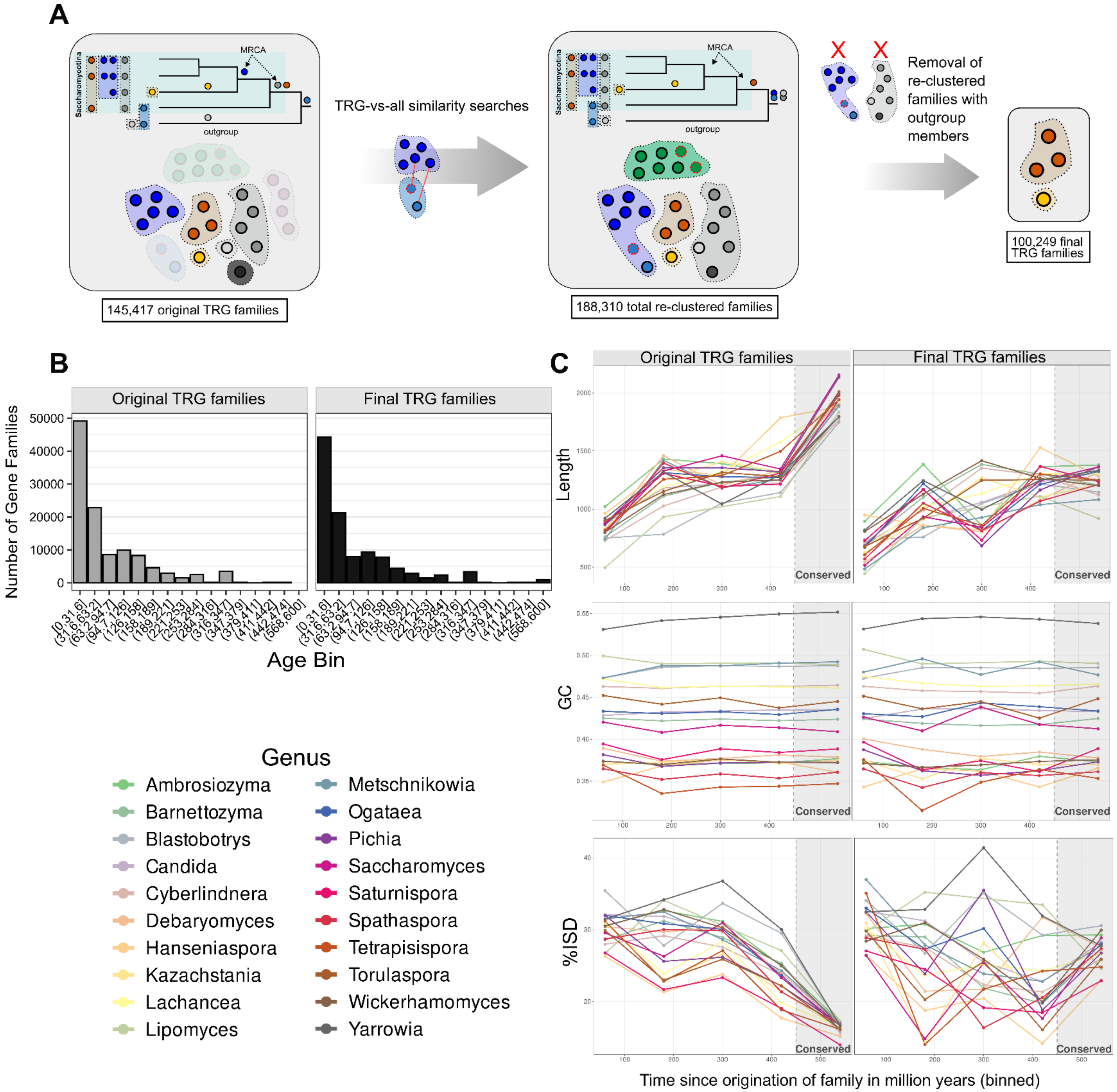


**Supp Figure 1 A**) Graphical representation of the re-clustering process and the change in the number of TRG families. **B**) Distribution of the number of TRG families per phylogenetic age bin before and after re-clustering. **C**) The evolutionary trajectories before and after re-clustering of top: CDS length, middle: GC content, bottom: intrinsic disorder of proteins. Each point corresponds to the average value for proteins of a given genus with an assigned age falling in a given bin.

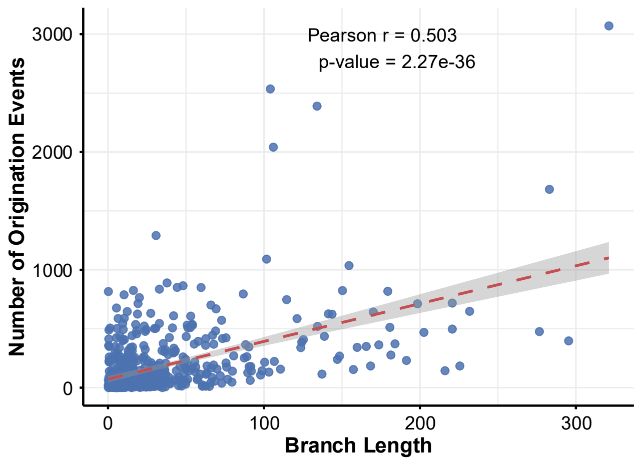


**Supp. Figure 2**: Correlation between phylogenetic branch length and the number of novel gene family origination events.

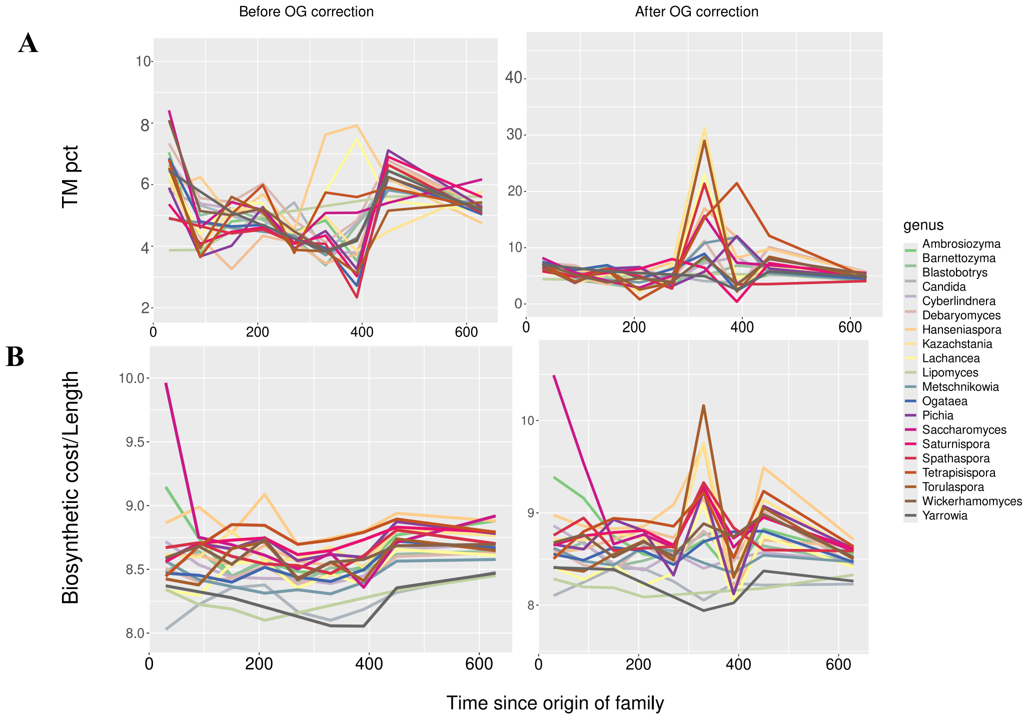


**Supp. Figure 3**: The property evolutionary trajectory of **A**) TM propensity, **B**) total biosynthetic cost normalized by protein length, before and after the gene family reassignment.

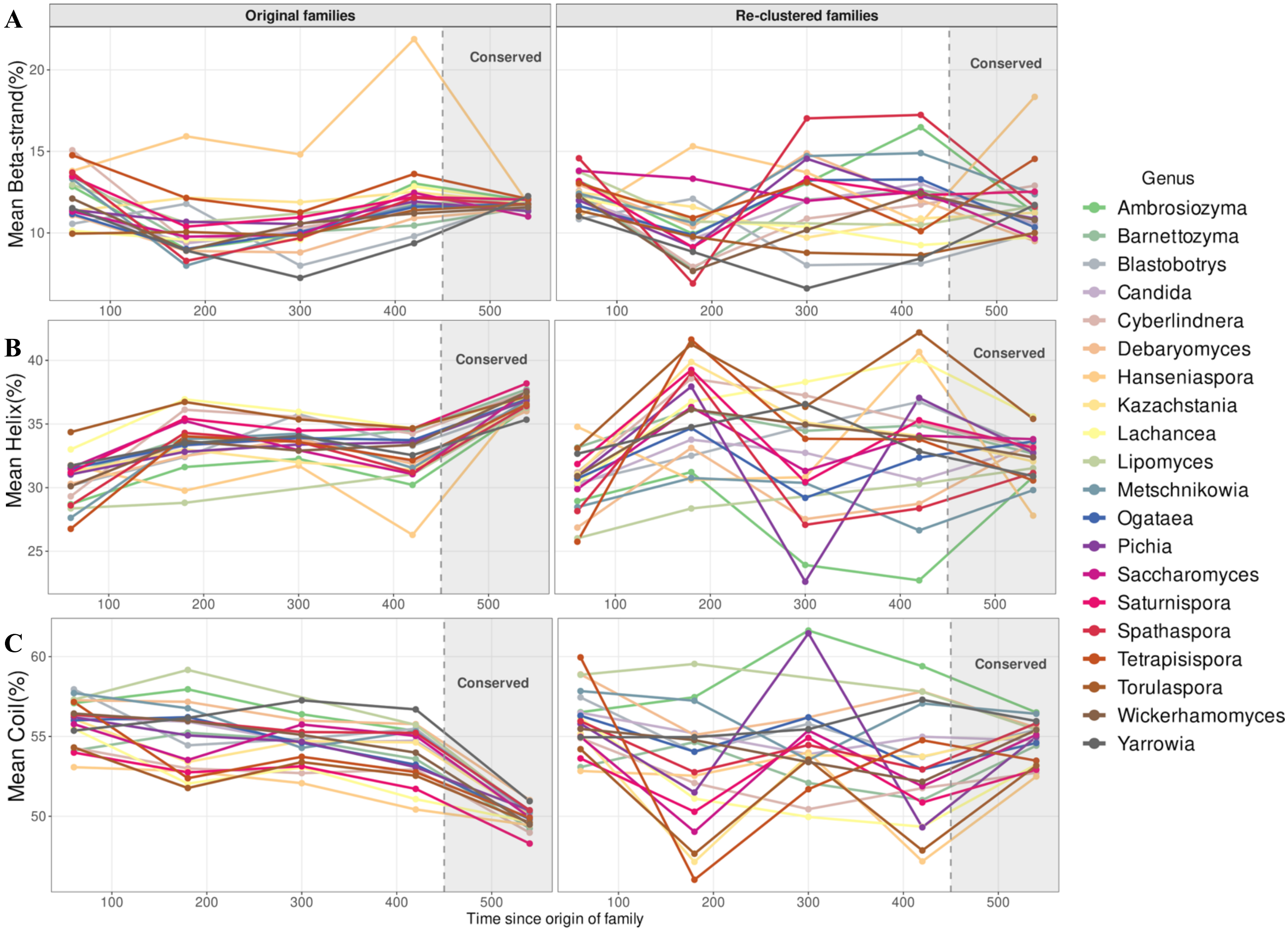


**Supp. Figure 4**: The property evolutionary trajectory of **A**) the average α-helices content of proteins **B**) the average β-strand content of proteins, **C**) the average coil content of proteins.

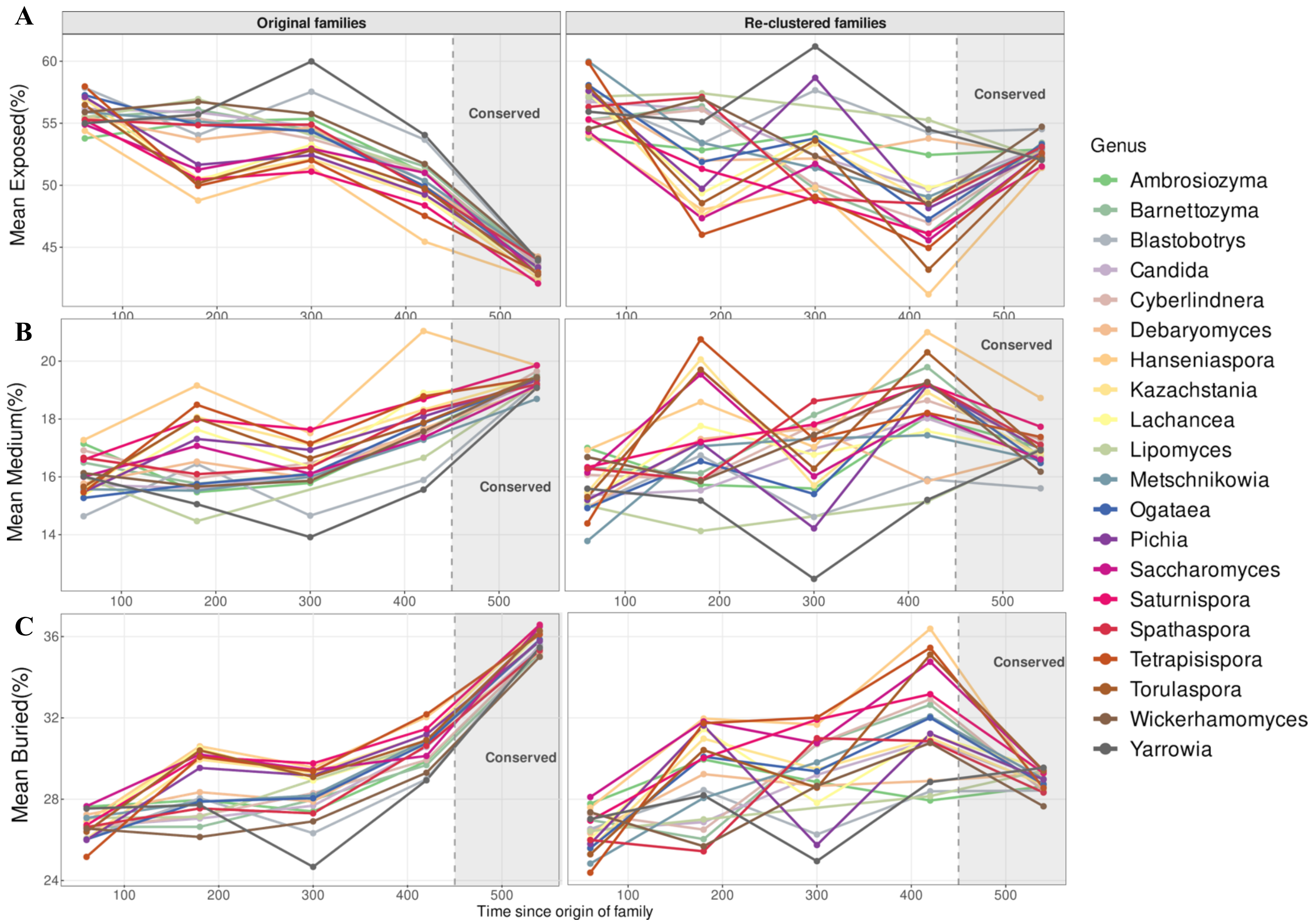


**Supp. Figure 5**: The property evolutionary trajectory of **A**) the percentage of the mean exposed amino acids per protein, **B**) the percentage of the mean partially exposed, partially buried amino acids per protein, **C**) the percentage of the mean buried amino acids per protein.

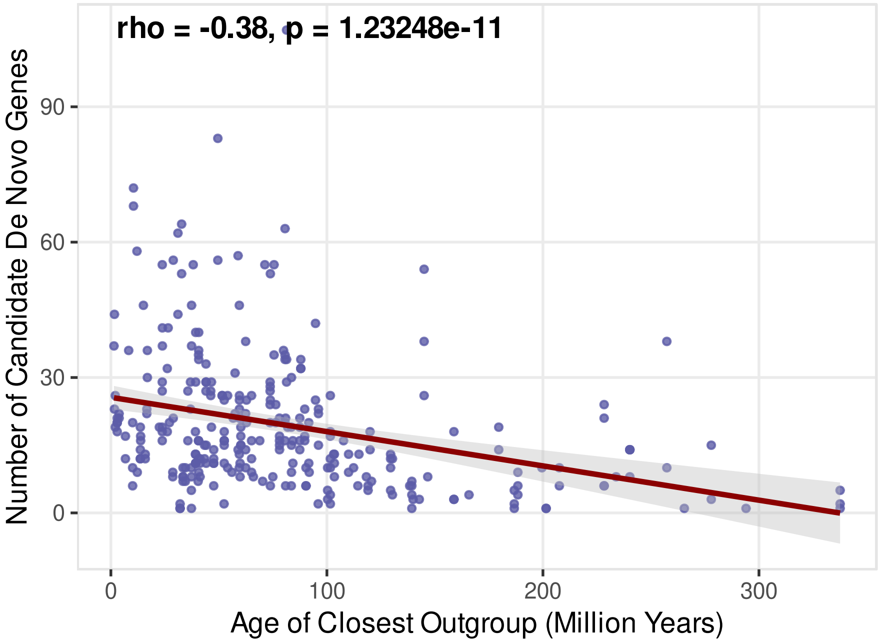


**Supp. Figure 6**: Correlation between the number of de novo genes in each species with the age of the furthest species used as outgroup.

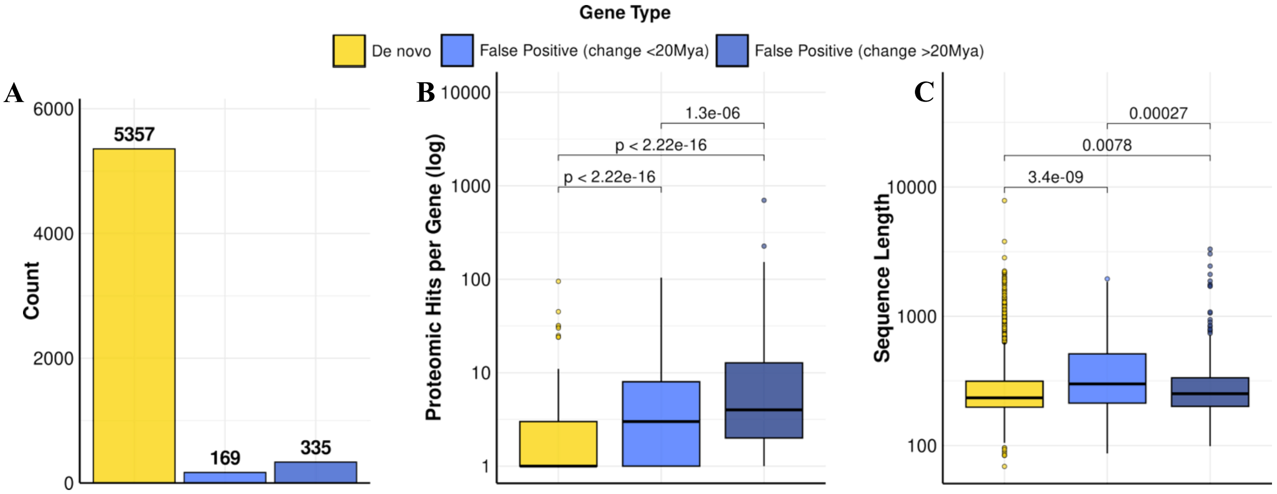


**Supp. Figure 7**: **A**) The number of de novo genes, FP and HGT cases, **B**) Comparison of the length between the true de novo genes, false positive de novo genes that the original age estimation was less than 20 my off, false positive de novo genes that the original age estimation was more than 20 Mya off and HGTs, **C**) Comparison of the number of proteomic hits between the true de novo genes, false positive de novo genes that the original age estimation was less than 20 my off, false positive de novo genes that the original age estimation was more than 20 my off and HGTs,

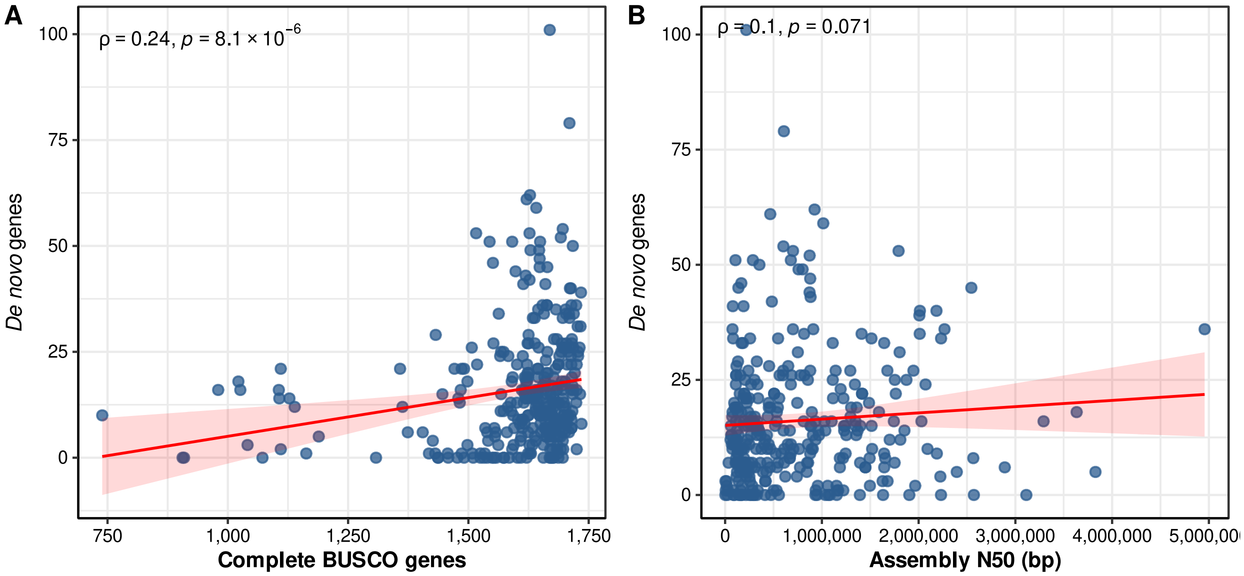


**Supp. Figure 8**: **A**) Correlation of the number of de novo genes in a species with the genome assembly quality of the species measured by the number of BUSCO genes present, **B**) Correlation of the number of de novo genes in a species with the genome assembly quality of the species measured by N50.

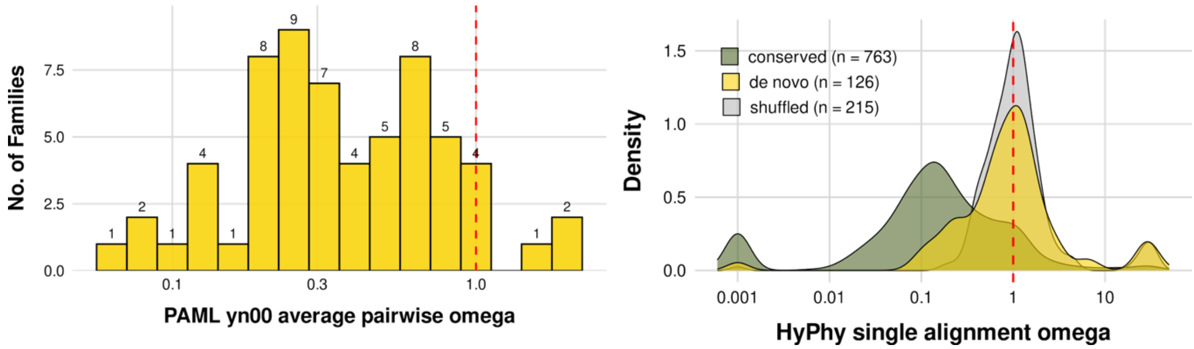


**Supp Figure 9**: **Left:** Distribution of $d_N/d_S$ values of 62 multimember, multispecies strict de novo gene families, as calculated using yn00 of PAML. **Right**:) Distribution of omega values (HyPhy) calculated from intra-specific alignments of strict de novo and conserved genes. Shuffled alignments of de novo genes are shown as negative controls.

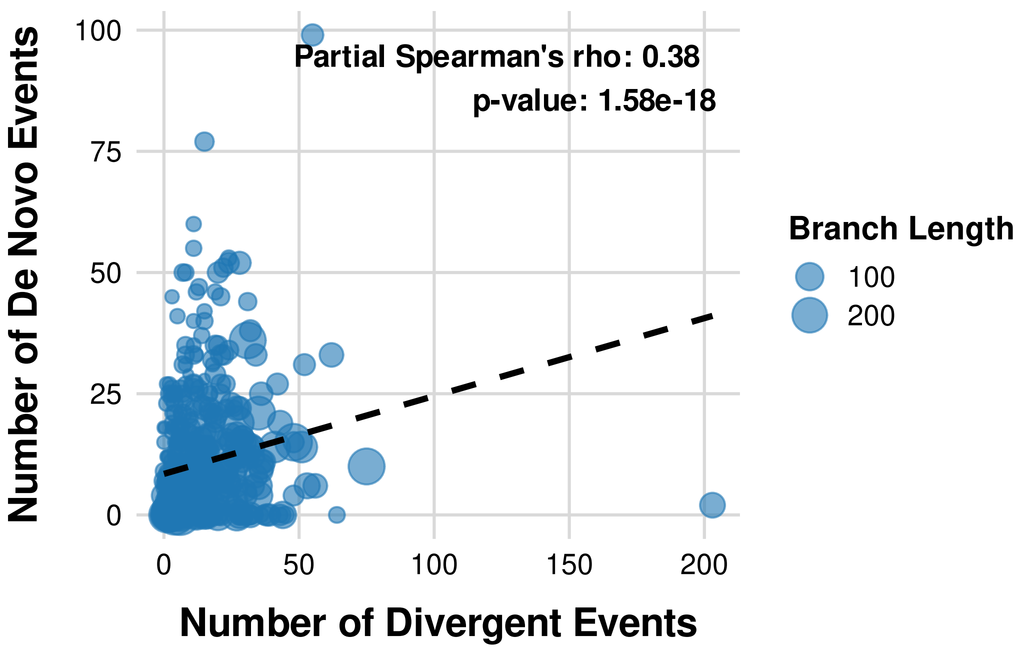


**Supp Figure 9**: Partial correlation between the de novo and divergent gene origination events with branch length.

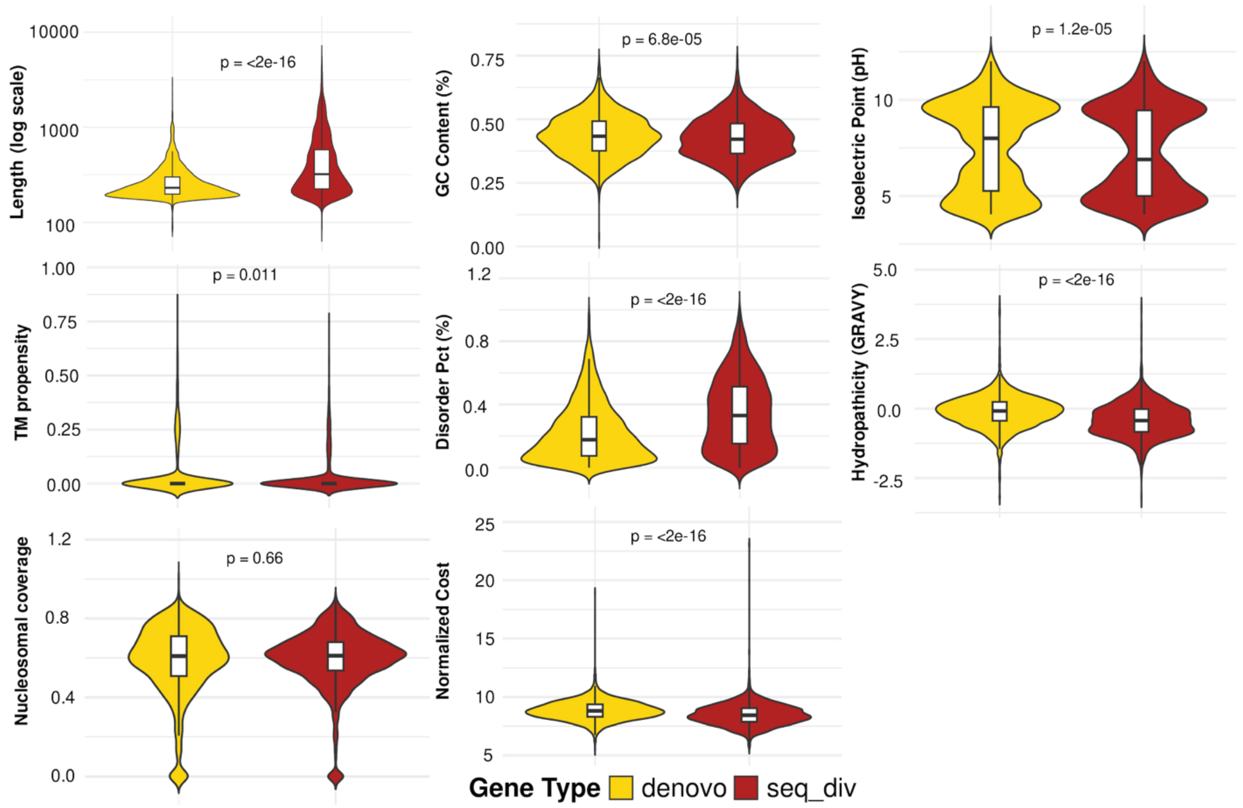


**Supp Figure 10**: Comparison of properties between young (<20 mya) de novo and divergent genes.

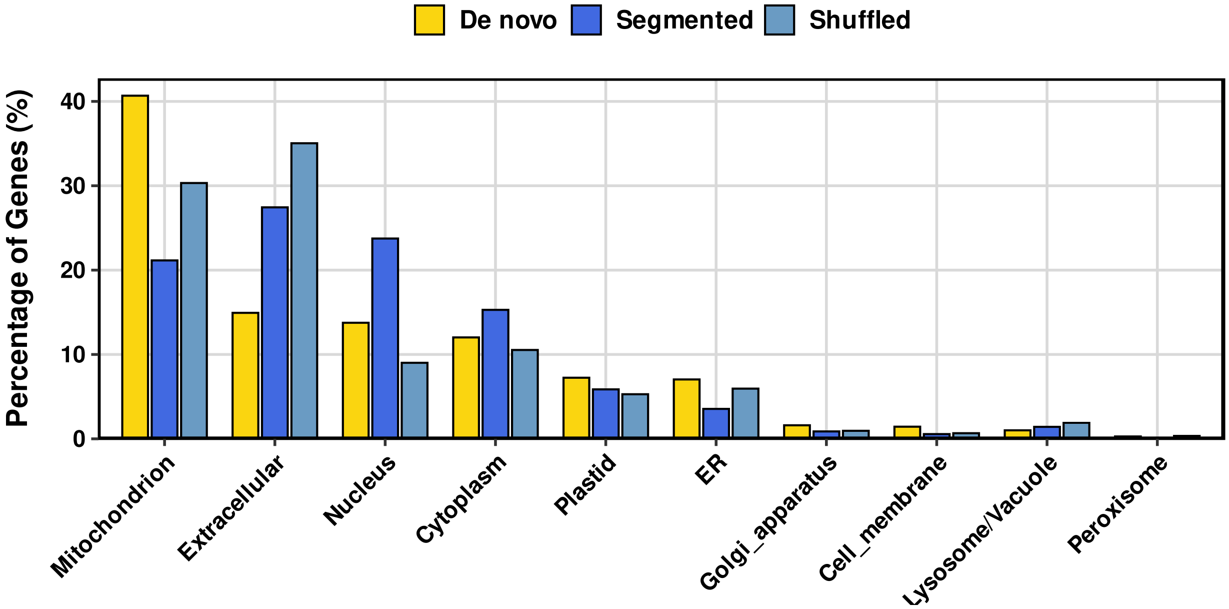


**Supp. Figure 11**: Subcellular localization predictions of de novo proteins and two control sets, one of shuffled de novo proteins and one with segments of conserved proteins of the same length as the de novo set, using DeepLoc1.

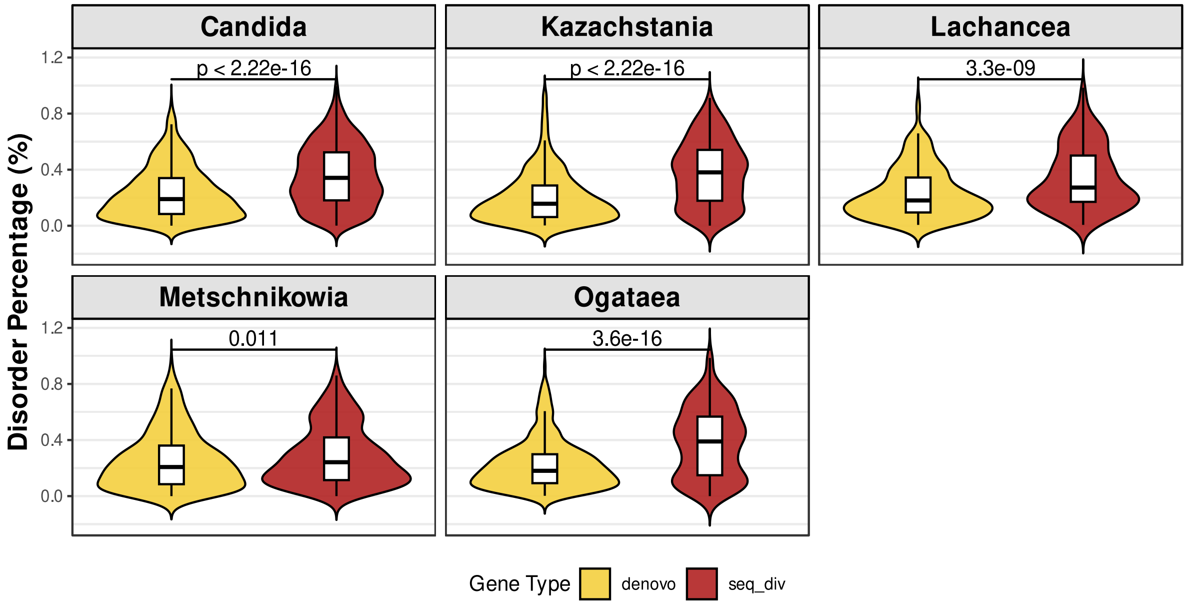


**Supp. Figure 12**: Comparison of the disorder percentage between of the de novo and divergent genes of the 5 most represented genera. Wilcoxon test P-values are shown.

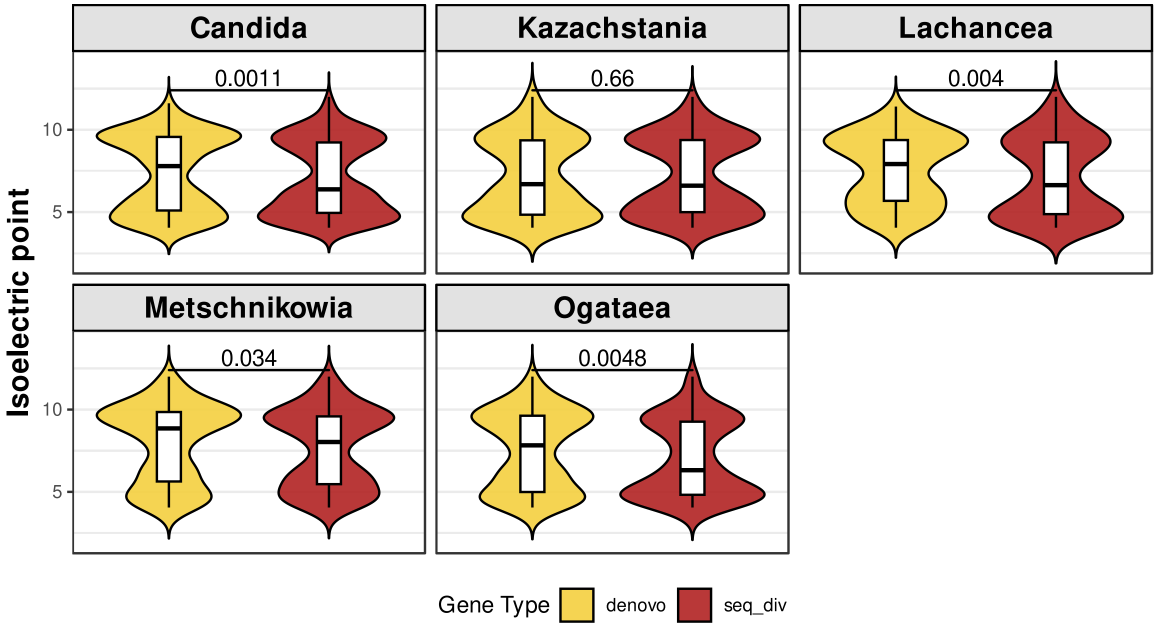


**Supp. Figure 13**: Comparison of the pI between of the de novo and divergent genes of the 5 most represented genera . Wilcoxon test P-values are shown.

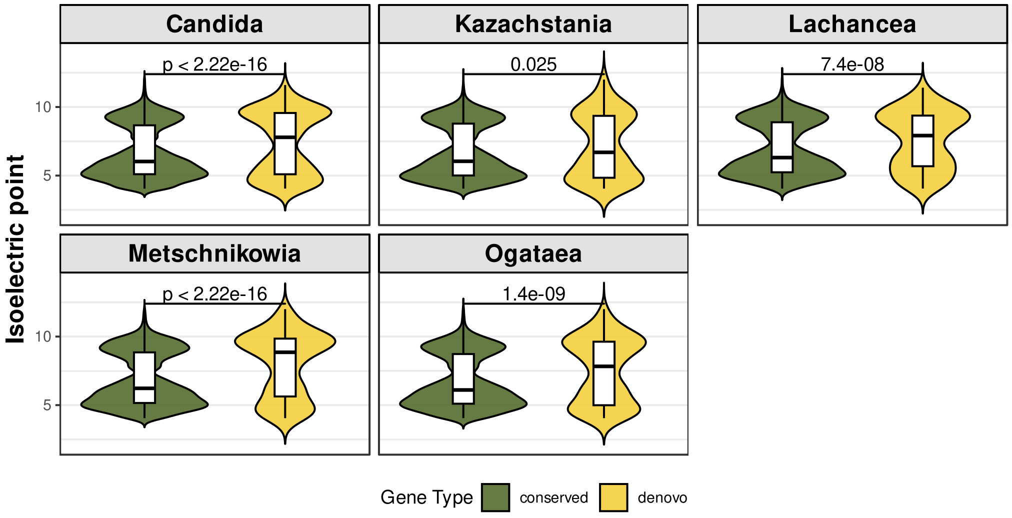


**Supp. Figure 14**: Comparison of the isoelectric point values between the de novo and conserved genes of the 5 most represented genera. Wilcoxon test P-values are shown.

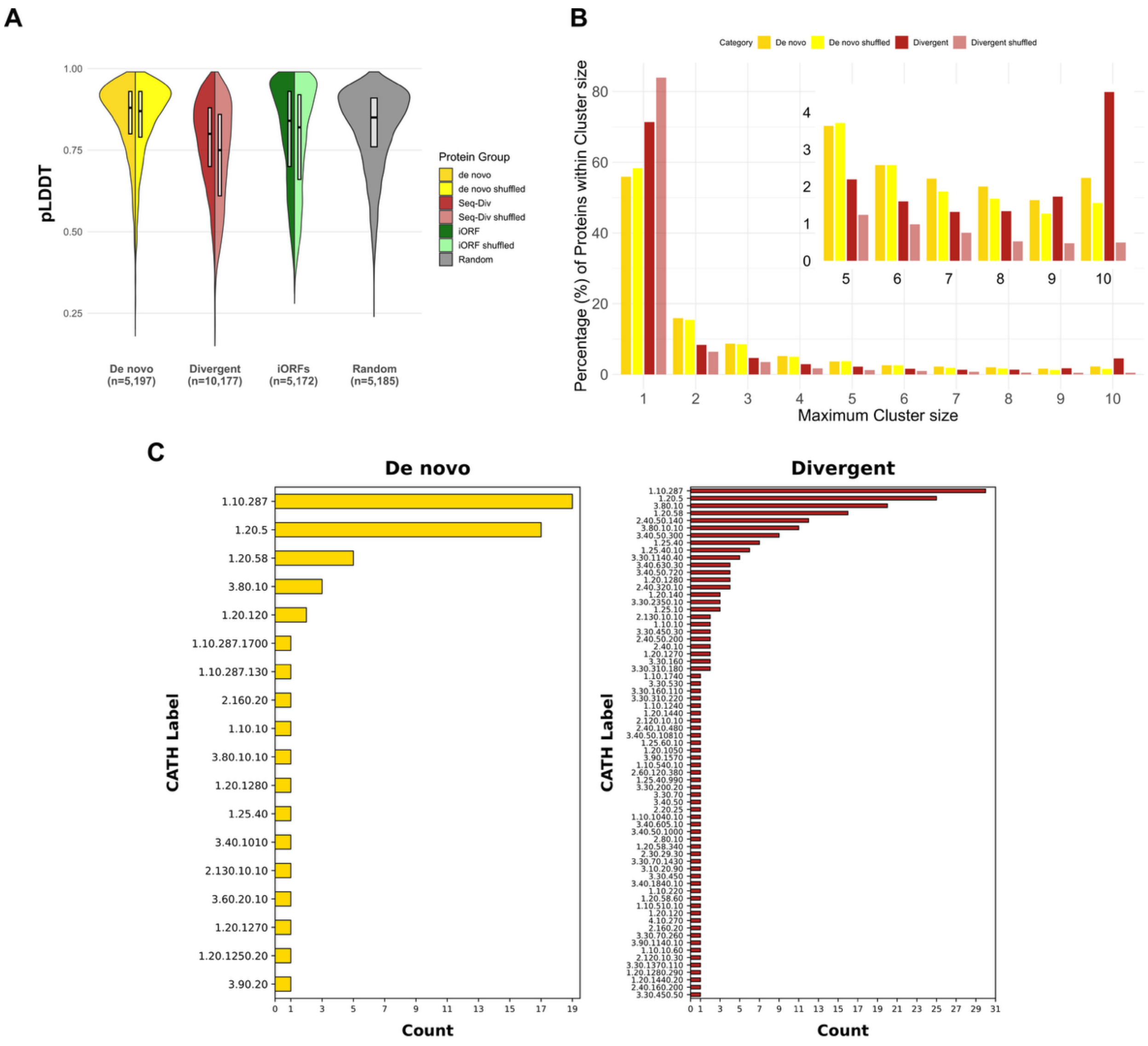


**Supp. Figure 15**: A) pLDDT value distributions for predicted structures of de novo, divergent, translated iORFs and random proteins, as well as shuffled controls. B) distribution of maximum cluster size within sets of 10 predicted structures resulting from 10 iterations for every de novo and divergent novel protein and associated shuffled controls. C) distributions of CATH hierarchy topologies and superfamilies to which belong TED domains matching to de novo and divergent HC structures.

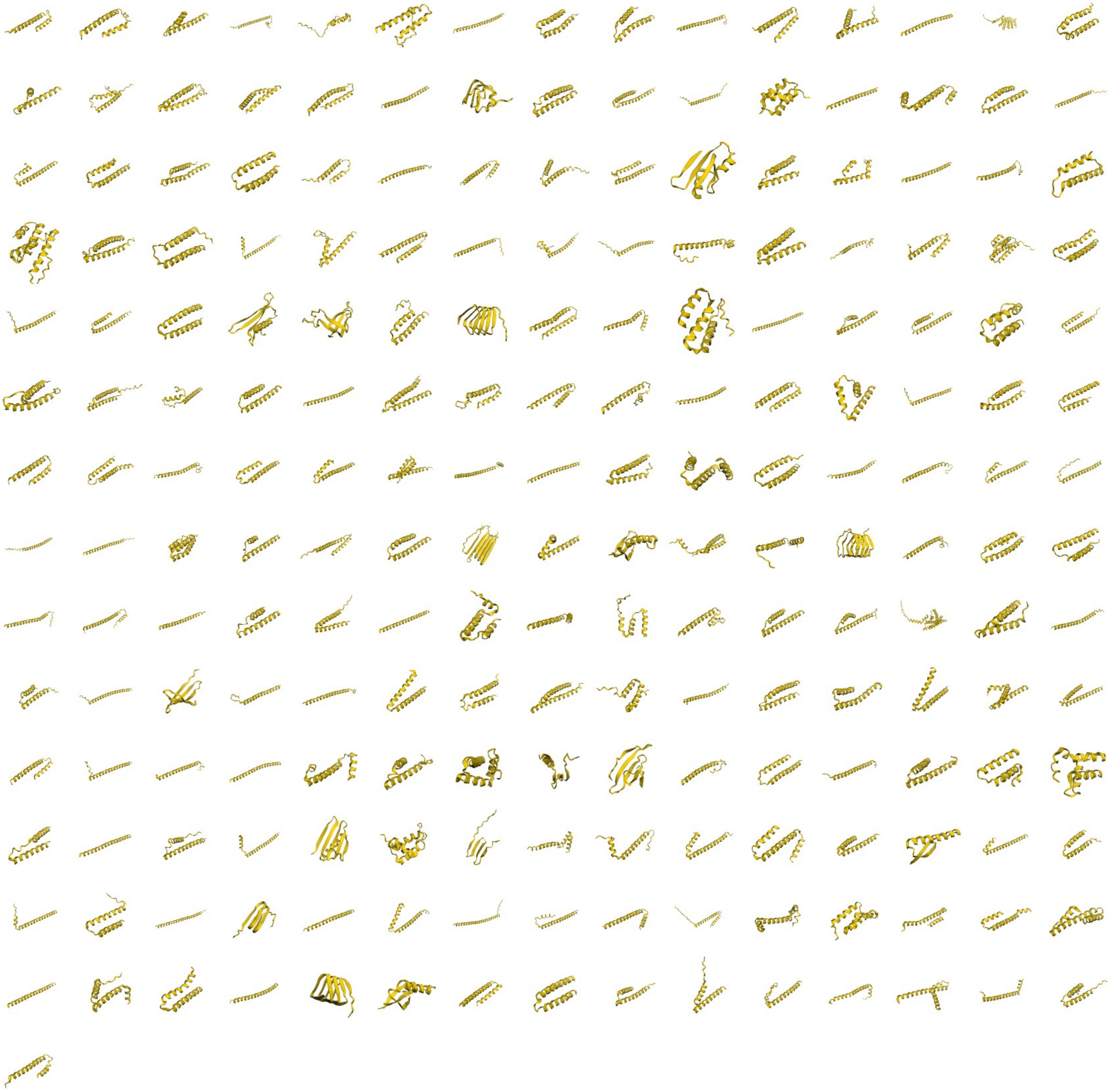

**Supp. Figure 16:** Novel high-confidence structural models of de novo proteins.

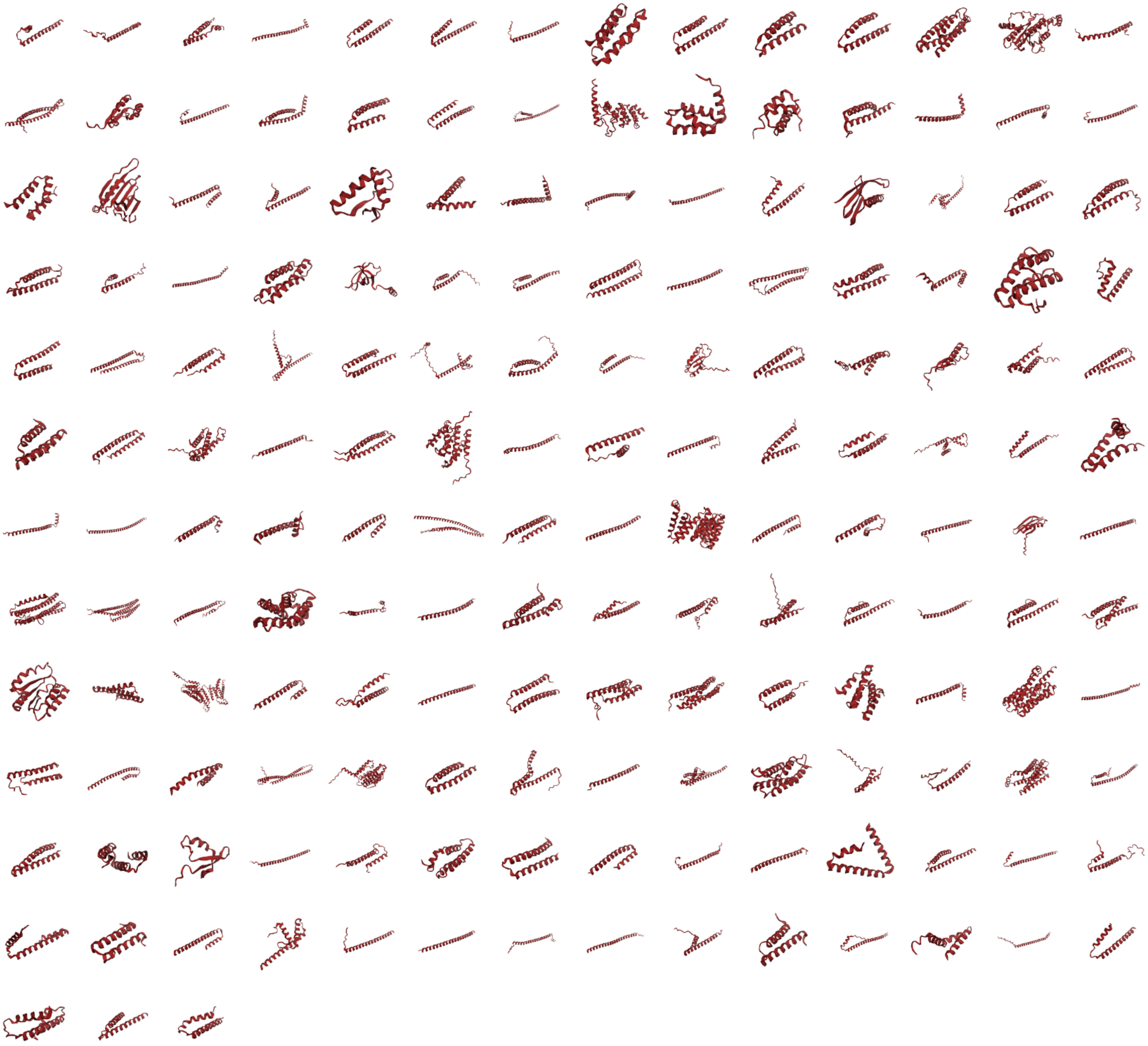

**Supp Figure 17:** Novel high-confidence structural models of divergent proteins.

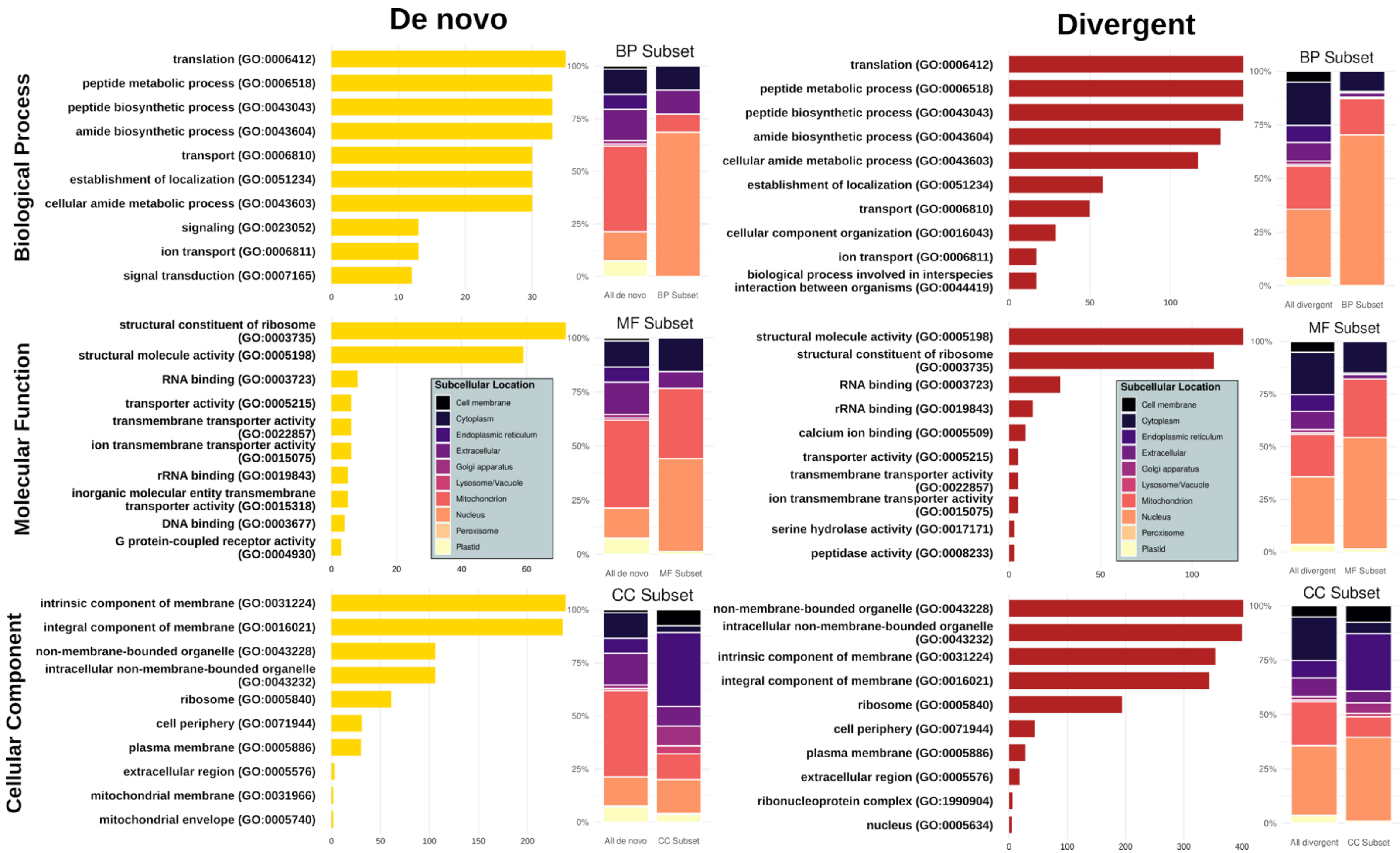


**Supp. Figure 18**: Top ten GO terms and associated protein counts, as predicted by DeepFRI for de novo and divergent proteins. Stacked barplots show percentages of predicted subcellular localization by DeepLoc for proteins assigned any of the top 3 GO terms on the right, as well as the background frequencies on the left (colour legends in the blue boxes).

### Supplementary Tables

**Supplementary Table 1**: Top Bacterial Matches for the HGT genes

**Supplementary Table 2**: Gene type lists (De novo, Divergent, HGT)

**Supplementary Table 3**: List of de novo genes in the strict set

**Supplementary Table 4**: List of de novo and divergent with robust structures

**Supplementary Table 5**: List of de novo and divergent genes with known structures within the structural databases AlphaFoldDB50.

**Supplementary Table 6**: List of de novo and divergent genes with known predicted domains

**Supplementary Table 7**: List of de novo and divergent genes with novel structures

**Supplementary Table 8**: Filtered list of de novo genes with novel structures

**Supplementary Table 9**: GO terms of de novo and divergent genes
